# Structure of LetB reveals a tunnel for lipid transport across the bacterial envelope

**DOI:** 10.1101/748145

**Authors:** Georgia L. Isom, Nicolas Coudray, Mark R. MacRae, Collin T. McManus, Damian C. Ekiert, Gira Bhabha

## Abstract

Gram-negative bacteria are surrounded by an outer membrane composed of phospholipids and lipopolysaccharide (LPS), which acts as a barrier to the environment and contributes to antibiotic resistance. While mechanisms of LPS transport have been well characterised, systems that translocate phospholipids across the periplasm, such as MCE (<u>M</u>ammalian <u>C</u>ell <u>E</u>ntry) transport systems, are less well understood. Here we show that *E. coli* MCE protein LetB (formerly YebT), forms a ∼0.6 megadalton complex in the periplasm. Our cryo-EM structure reveals that LetB consists of a stack of seven modular rings, creating a long hydrophobic tunnel through the centre of the complex. LetB is sufficiently large to span the gap between the inner and outer membranes, and mutations that shorten the tunnel abolish function. Lipids bind inside the tunnel, suggesting that it functions as a pathway for lipid transport. Cryo-EM structures in the open and closed states reveal a dynamic tunnel lining, with implications for gating or substrate translocation. Together, our results support a model in which LetB establishes a physical link between the bacterial inner and outer membranes, and creates a hydrophobic pathway for the translocation of lipids across the periplasm, to maintain the integrity of the outer membrane permeability barrier.

---

The bacterial outer membrane (OM) constitutes a formidable barrier that protects the cell against antibiotics and other harsh environmental stresses. Understanding the machinery involved in building and maintaining the OM is critical for the development of new antimicrobials targeting these fundamental, conserved pathways. In Gram-negative bacteria, the OM is an asymmetric lipid bilayer, with a phospholipid inner leaflet and a lipopolysaccharide (LPS) outer leaflet, and this asymmetry is critical for its barrier function. To generate this barrier, lipids must be transported across the cell envelope. Membrane lipids are poorly soluble in water, and must be protected from solvent during transport across the hydrophilic periplasmic space between the inner membrane (IM) and OM. A series of studies have unraveled how the LPS transport system employs an ABC transporter in the IM to drive movement of LPS molecules across a periplasmic bridge to an OM complex that mediates their insertion into the outer leaflet^1–5^. Analogous systems likely exist for the export of nascent OM phospholipids, and for lipid import, as in the case of exogenous or mislocalised lipids, but the identities and mechanisms of these systems are not well established. MCE transporters have recently emerged as candidates for lipid translocation across the periplasm^6–8^.

MCE proteins (which contain <u>M</u>ammalian <u>C</u>ell <u>E</u>ntry domains) have been shown to interact with IM ABC transporters, or are present in operons with integral membrane proteins that have been proposed to encode membrane transporters^7,9,10^. These multi-subunit MCE systems are broadly conserved in double-membraned bacteria and chloroplasts^11^, and have been implicated in the import and/or export of lipids, cholesterol or other hydrophobic molecules^6,8,12^. Members of this transporter family play an important role in the outer membranes of both Gram-negative bacteria^6,9^ and *Mycobacteria*^*13*^, and are critical during host infection of various bacterial pathogens^14,15^.

The best characterised MCE transport system, called Mla, has been implicated in the <u>M</u>aintenance of OM <u>L</u>ipid <u>A</u>symmetry in *E. coli* by importing mislocalised phospholipids from the OM to the IM^6^, and consists of three main parts: 1) an OM complex, MlaA-OmpC, that may mediate the extraction of lipids from the OM, 2) a lipid carrier protein, MlaC, that ferries lipids across the periplasm, and 3) an IM ABC transporter complex, MlaFEDB, that may mediate the insertion of lipids into the IM. More recent work has suggested, however, that the Mla system may in fact drive lipid export, wherein MlaFEDB may use ATP hydrolysis to extract lipids from the IM and MlaA-OmpC may mediate their insertion in the OM^16,17^. Thus, the directionality of the Mla phospholipid transporter and potentially other MCE protein family members remains an open question.

In the MlaFEDB complex, the MCE subunit MlaD forms a homohexameric ring on the periplasmic side of the IM. A hydrophobic tunnel through the centre of the MCE ring was proposed to form part of the lipid translocation pathway^7^, while a soluble protein, MlaC, shuttles lipids across the remaining periplasmic gap. However, other *E. coli* MCE genes, such as *pqiB* are much larger than *mlaD* and encode proteins with multiple MCE domains that can create much longer architectures^7^. Here, we report the high-resolution cryo-electron microscopy (cryo-EM) structure of one of the largest MCE proteins identified to date, a protein from *E. coli* that we name LetB for <u>L</u>ipophilic <u>E</u>nvelope-spanning <u>T</u>unnel B (formerly YebT). We show that LetB forms a tunnel sufficiently long to mediate lipid transport directly between the IM and OM without the need for a shuttle protein. Thus, like LPS, phospholipids may also be transported across the periplasm by large transenvelope complexes.

## Results

### LetB forms a periplasm-spanning tunnel

*LetB* from *E. coli* is a large MCE gene encoded in an operon together with *letA* (formerly *yebS*), an integral inner membrane protein (**Fig. 1a**). *E. coli* and Salmonella strains lacking *letAB* exhibit phenotypes consistent with outer membrane defects^9,15^ **(Extended Data Fig. 1**). *LetB* encodes seven MCE domains in a single polypeptide chain (MCE1 - MCE7) with an N-terminal transmembrane helix. Our previous low resolution negative stain EM data suggested that LetB resembles an elongated, segmented rod^7^. To gain an understanding of how LetB may function to modulate OM integrity, we used cryo-EM to determine a high-resolution structure of LetB (residues 43-877, **Fig. 1a**). We collected data on a Krios microscope equipped with a K2 camera (**Extended Data Table. 1**), which we processed in Relion^18^. 2D class averages clearly show the overall arrangement of LetB, which is an elongated, hollow, hexameric structure with seven layers (**Fig. 1b**). Extensive 3D classification, signal subtraction, and refinement schemes yielded a set of 12 final maps covering various subvolumes (**Extended Data Figs. 2 and 3a, b**). The resolution of these maps ranged from ∼3 - 4 Å, allowing us to build a complete structural model of LetB (**Extended Data Table 1, Extended Data Fig. 3c**).

**Figure 1.**
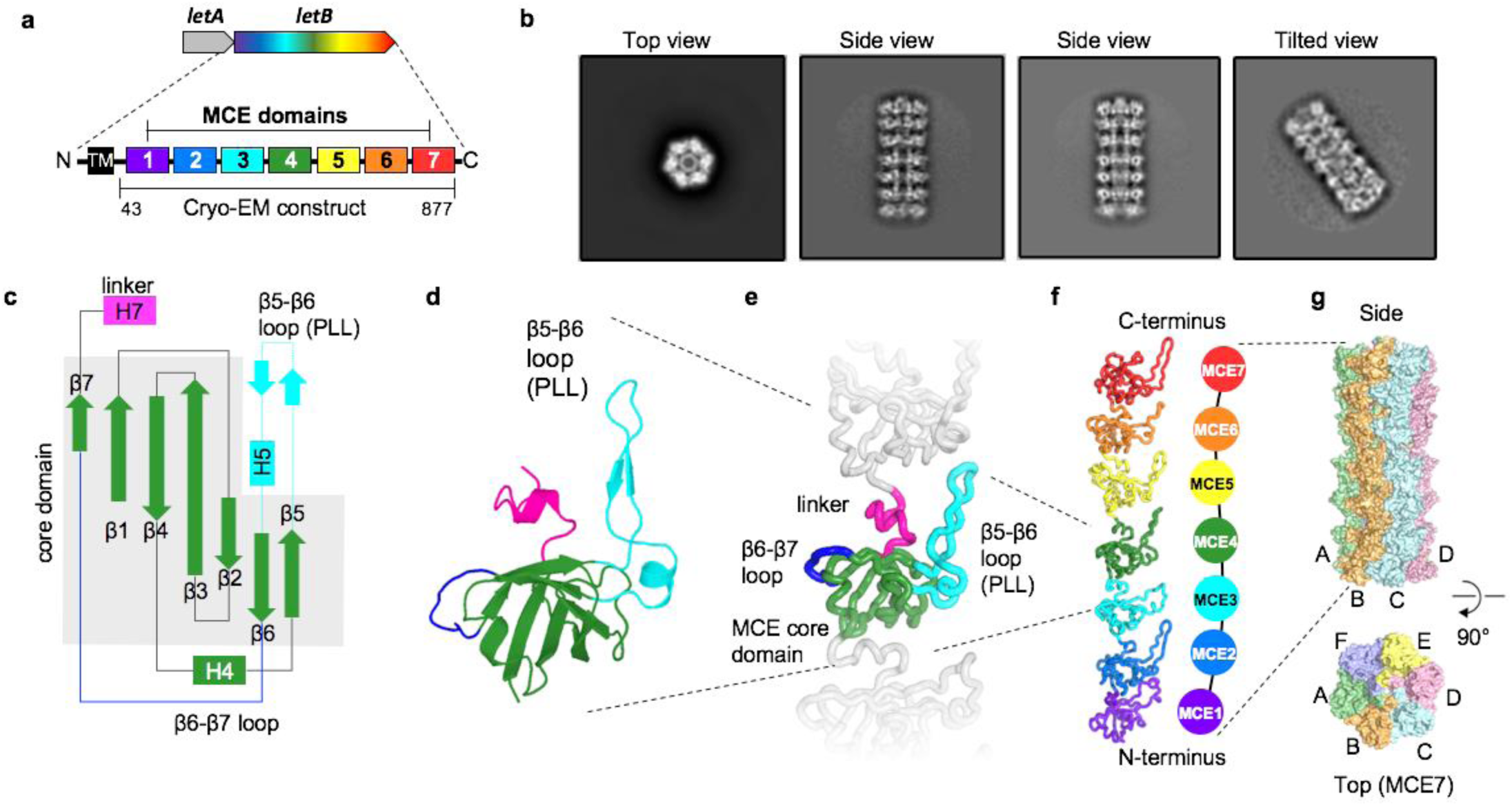
Cryo-EM structure of LetB. **a**. *letAB* operon and schematic of LetB domain organisation. The single-pass transmembrane helix (TM) is shown in black and MCE domains 1-7 in colours. The TM was deleted in the construct used for cryo-EM. **b**. Representative 2D class averages of LetB cryo-EM data. **c.** Topology of an MCE domain from LetB. β-strands and α-helices are depicted as arrows and rectangles, respectively. **d**. Cartoon representation of a representative MCE domain (MCE4) from LetB. Colouring of secondary structure elements corresponds to **c. e**. MCE4 domain (coloured) shown in the context of the preceding and following MCE domains (grey). Each domain is composed of a β-barrel core (green). Loops that vary significantly between domains are: β6-β7 loop (blue), and β5-β6 loop (cyan), which we term pore-lining loop (PLL), see **Fig. 2c**. A short helical linker (magenta) connects one domain to the next. **f**. Cartoon representation LetB protomer showing the linear arrangement of seven MCE domains, from the N-terminal MCE1 to the C-terminal MCE7. **g.** Surface representation of LetB. Six protomers (A-F) oligomerise laterally to form a homohexameric assembly. Each protomer runs the length of the LetB assembly.

The sequences of the seven MCE domains in LetB differ considerably, with sequence identity ranging from 17% to 28% in pairwise comparisons, and the domains of LetB are also highly diverged from the other *E. coli* MCE proteins MlaD and PqiB (**Extended Data Fig 4a,b**). However, each MCE domain has a structurally conserved core, consisting of a seven stranded β-barrel (**Fig. 1c,d**), similar to the MCE domains of PqiB and MlaD^7^. In addition, we identify two main regions that are variable between the structures of MCE domains (**Fig. 1e**). The first variable region is the β6-β7 loop on the exterior surface of the β-barrel, which ranges from 9 to 15 residues in length and in which only MCE5 contains a helix. The second variable region is the β5-β6 loop, which ranges from 17 to 27 residues, with an average length of 20 residues in LetB and consists of a β-hairpin followed by a short ∼two-turn helix.

The seven MCE domains of a single protomer of LetB resemble seven beads on a string (**Fig. 1f**), with the order of the domains in the primary sequence determining the order of MCE “beads” on the string. The seven MCE domains are stacked almost directly on top of one another, and each domain is connected to the next by a short helical linker. Six copies of this linearly configured LetB protomer associate laterally to form a homohexameric assembly (**Fig. 1g**). In the assembly, each MCE domain associates with the five other equivalent MCE domains from adjacent protomers to form a ring. The seven MCE domains lead to a final assembly of seven distinct rings, with a ring of MCE1 domains at the N-terminus and the MCE7 ring at the C-terminus (**Fig. 2a**). These seven MCE rings that make up LetB stack to form an elongated macromolecular barrel with a single protomer extending the entire length of the barrel. The N-terminal transmembrane helix of each protomer is expected to be anchored in the IM, as previously reported^9^, with the MCE1 ring abutting the IM and the MCE7 ring closest to the OM.

**Figure 2.**
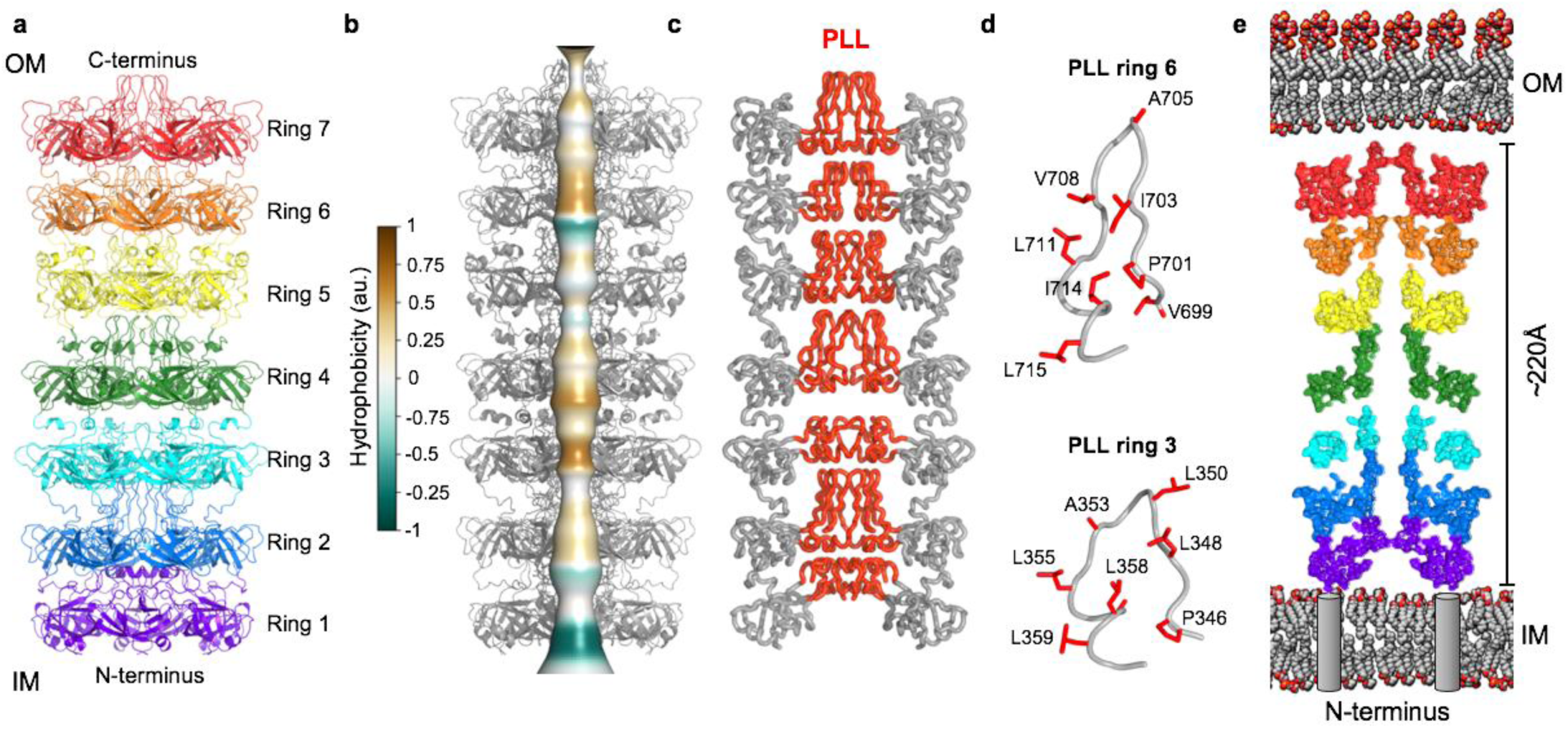
LetB hexamers form a tunnel. **a.** Cartoon representation of a LetB hexamer showing 7 stacked rings, each formed from the association of 6 identical MCE domains. **b.** Composite model of LetB in the open state (further described in Fig. 4 and associated text), showing tunnel running through the protein assembly. Tunnel is depicted as smooth surface coloured by the hydrophobicity of pore-facing residues, calculated using CHAP^<u>54</u>^. **c**. Ribbon representation of LetB. Pore-lining loops (PLLs; red) from each MCE domain form the central tunnel. **d**. Representative PLLs (PLL3 and PLL6) with pore facing residues shown as red sticks. Most pore-facing residues are hydrophobic. **e**. Cross-section of the LetB tunnel oriented in the context of the IM and OM, with the N-terminus anchoring the complex in the IM and the C-terminus towards the OM. Grey cylinders represent transmembrane helices (TM, not present in the structure) at their expected positions. The width depicted between the IM and OM is within a range of 210 −240 Å, based on previously measured periplasmic widths, 210 Å^<u>22</u>^ and 230 Å^<u>21</u>^, and deduced from the periplasmic spanning region of AcrAB-TolC, 240 Å^<u>59</u>^.

A continuous central tunnel runs through LetB, which is largely hydrophobic (**Fig. 2b,c**), similar to that of the other two *E. coli* MCE proteins, PqiB and MlaD^7^. Though evolutionarily unrelated, the multidrug efflux pump AcrAB-TolC forms a similarly elongated assembly in the periplasm, and uses proton motive force to transport substrates through its central tunnel out of the cell (**Extended Data Fig. 5**). The average diameter of the LetB tunnel is ∼14 Å, narrower than the tunnel running through AcrAB-TolC, but large enough for the passage of lipids and other small molecules. While in AcrAB-TolC, the tunnel is formed via the tight association of helices to form a rigid barrel, in LetB the tunnel is largely formed by the conformationally dynamic β5-β6 loops that emerge from each MCE domain, which we term pore-lining loops (PLLs) (**Fig. 2d**). The length of LetB is ∼220 Å, comparable to the width of the periplasmic space^19–22^ (**Fig. 2e**) and to the length of the periplasm spanning region of AcrAB-TolC^23,24^. In summary, LetB is poised to serve as a tunnel for the transport of small molecules between the IM and OM by creating a bridge across the periplasm, perhaps via interactions with LetA in the IM and proteins or lipids in the OM.

### Number of MCE domains determines tunnel length

The structure of LetB implies that MCE domains function as ring-shaped modular building blocks. This led us to hypothesise that in multi-domain MCE proteins, the number of MCE domains in the primary sequence specifies the number of rings stacked in the final assembly. Indeed, the primary sequences of *E. coli* MlaD, PqiB, and LetB consist of one, three, and seven tandem MCE domains, respectively, and structures of these three proteins revealed that they consist of one, three, and seven stacked rings (**Fig. 3a**). To further test this hypothesis, we selected MCE proteins from other bacterial species that were predicted to contain four, five, six or eight MCE domains^9^ (**Extended Data Fig. 6a**), and visualised them by negative stain EM. 2D class averages for the proteins containing four, five, and six tandem MCE domains clearly revealed multiple layers of density, consistent with barrel structures assembled from four, five, and six stacked rings, respectively. The eight domain MCE protein was less stable, often resulting in smaller, likely monomeric, particles; when oligomers were formed, these contained eight stacked rings (**Fig. 3b, Extended Data Fig. 6b, Supplementary Data Fig. 2a,b**). 2D class averages of the top view of all proteins revealed that they are approximately 6-fold symmetric like LetB and other MCE proteins, suggesting that they also assemble to form homo-hexamers (**Extended Data Fig. 6b**). Overall, these data indicate the MCE domains are modular building blocks of larger proteins, with each domain in the primary sequence specifying one ring in the resulting structure. From the 2D classes of these proteins and our structures of MlaD (1 ring), PqiB (3 rings), and LetB (7 rings), we estimated the lengths of the one, three, four, five, six, seven, and eight ring MCE barrels to be ∼25, ∼95, ∼140, ∼165, ∼190, ∼220, and ∼260 Å, respectively. Thus, naturally occurring MCE proteins have evolved to form barrels with a range of lengths, perhaps in part to adapt to differences in the mean periplasmic width in different species and to accommodate larger or smaller interacting complexes in the IM or OM.

**Figure 3.**
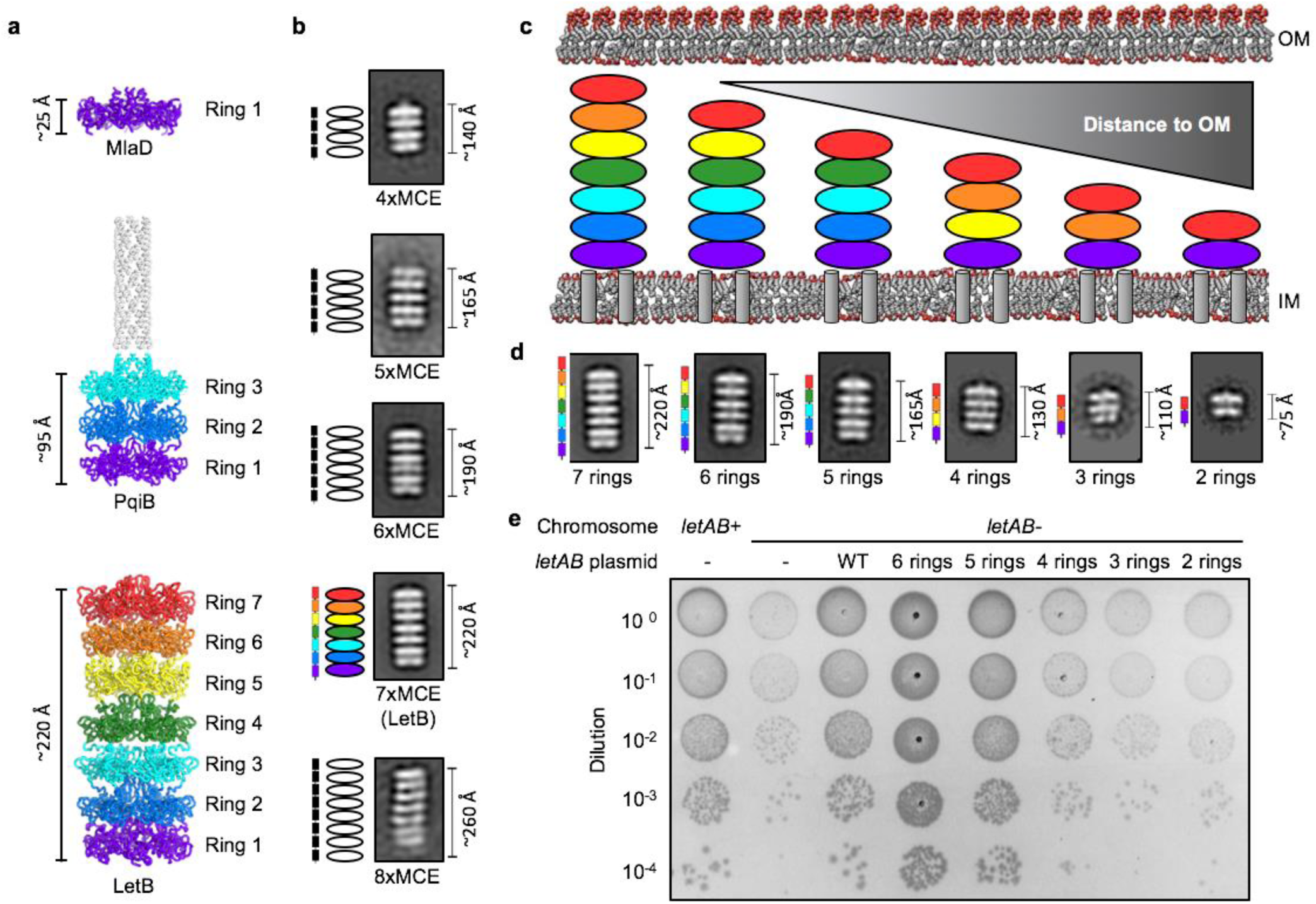
MCE domains are modular building blocks that determine the length of MCE protein assemblies. **a.** Cartoon representations of *E. coli* MCE proteins MlaD (PDB ID: 5UW2), PqiB (PDB ID: 5UVN) and LetB, which contain one, three and seven MCE domains, respectively, and form architectures with the corresponding number of rings. **b**. Negative stain EM 2D class averages of naturally occurring proteins encoding four, five, six, seven (LetB), and eight MCE domains (**Extended Data Fig. 6**). Schematics to the left of 2D class averages indicate number of MCE domains present in the gene (rectangles) and expected number of rings based on modularity of MCE domains (ovals) **c.** Truncated constructs designed to modulate number of LetB rings and their expected length in the context of the periplasm. As number of LetB domains is reduced, distance to OM is predicted to increase. Colour coding is the same as in **a**; see **Methods** for rationale of construct design. **d.** Negative stain EM 2D class averages of LetB truncation mutants depicted in **c**, but with the TM helix deleted for each mutant. Domain organisation of the gene is shown to the left of the 2D class average. The 7 ring 2D class average is the same as the 7xMCE class in (**b**). **e.** Cellular assay for the function of LetB truncation mutants depicted in **c.** 10-fold serial dilutions of the indicated cultures spotted on plates containing LSB and incubated overnight. The *letAB* double mutant grows poorly in the presence of LSB, but can be rescued by the expression of *letAB* constructs containing *letB* encoding five, six, or seven (WT) MCE rings. Expression of constructs with *letB* containing two, three, or four rings fail to rescue.

### Length of LetB affects its function *in vivo*

Given that the length of MCE proteins varies with the number of MCE domains, and the length of LetB is approximately the same as the reported width of the periplasm in *E. coli*^*19–22*^, we hypothesised that LetB may physically bridge the gap between the IM and OM, creating a continuous path for lipid transport. If this model is correct, then the length of LetB would be important for its function. To test this hypothesis, we engineered shorter mutant forms of LetB, which contained two, three, four, five or six MCE domains instead of seven, as in the wild-type protein (**Fig. 3c**, see **Methods** for rationale of mutant design). To assess if these engineered mutants were folded and assembled into shorter barrels of the expected size, we overexpressed and purified each mutant, and imaged the proteins by negative stain EM. 2D classification revealed that all of the LetB truncation mutants folded and homo-hexamerised to form the expected number of rings (**Fig. 3d**).

To assay the function of these shorter LetB variants, we took advantage of a genetic interaction between *pqiAB* and *letAB* mutants. Disruption of *pqiAB* results in a moderate growth defect in the presence of the detergent lauryl sulfobetaine (LSB), while the growth of a *letAB* mutant is unaffected^9^. However, disruption of *letAB* in combination with *pqiAB* (*pqiAB letAB* double knockout) significantly exacerbates LSB sensitivity, and complementation with wild-type *letAB* from a plasmid relieves this exacerbation, as previously described^9^ (**Extended Data Fig. 1c**). We used this complementation assay to assess whether or not the shorter LetB variants in a plasmid can also relieve the exacerbated phenotype of the *pqiAB letAB* double knockout. LetB variants that are two, three and four rings in length were unable to restore growth of the *pqiAB letAB* mutant in the presence of LSB, indicating that these proteins are non-functional (**Fig. 3e**), despite being stable and properly folded *in vitro* (**Fig. 3d**). In contrast, LetB variants that were five and six rings in length did restore growth of the *pqiAB letAB* mutant in the presence of LSB, indicating that they function as well as the wild-type seven-ring protein (**Fig. 3e**). These results are consistent with the idea that the length of LetB may be important for its function, perhaps reflecting a minimum length required to bridge the distance between the IM and OM.

### Conformational dynamics modulate tunnel diameter

Our cryo-EM density map showed that the local resolution varied considerably between different regions of LetB: while rings 2-4 were at the highest resolution and well-ordered, rings 1, 5, 6, and 7 were at a much lower resolution (**Fig. 4a, Extended Data Fig. 3a**). One possibility was that conformational heterogeneity was leading to lower resolution. To tease apart different conformations, we explored multi-body refinement^25^ and 3D variability analysis^26^, which showed the flexibility of the overall structure (**Supplementary Video 1**). Using several masking and signal subtraction strategies (**Extended Data Fig. 2**), we identified 8 classes ranging from 2.96 Å to 3.78 Å resolution, capturing the conformational dynamics of the sample. Rings 1, 5, 6 and 7 show multiple discrete conformational states, while rings 2, 3 and 4 are relatively rigid and are resolved in one conformation (**Fig. 4b, Supplementary Video 2, Extended Data Fig. 2** and **3a**). We focus our analysis on the two most distinct conformational states, in which the central tunnel is either open or closed.

**Figure 4.**
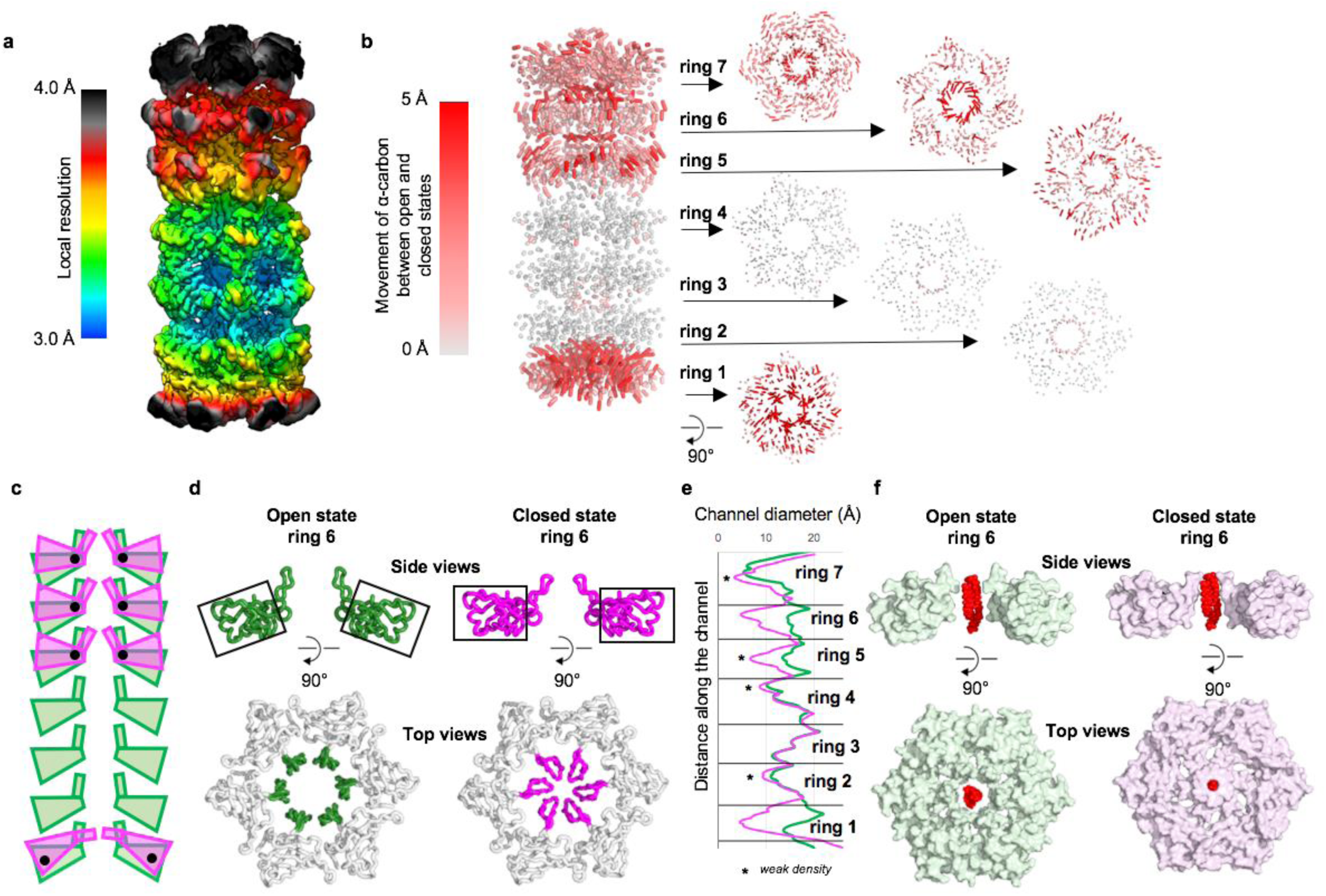
Conformational dynamics showing open and closed states of LetB. **a.** Local resolution of the LetB density map before 3D classification and local refinement. **b**. Movement of the α-carbons between the composite models of the closed and open states. Red trajectories indicate regions of highest displacement. Larger movements are primarily seen for rings 1, 5, 6 and 7, but not rings 2-4. Side view of intact LetB is shown at left. Arrows link side view to top view of the corresponding ring. **c**. Schematic representation of domain and PLL movements observed in rings 1, 5, 6 and 7 from the open (green) to the closed (magenta) states, with axis of rotation represented as black dots. **d**. The orientation of the domains (top) and PLLs (bottom) for ring 6 in open (green) and closed (magenta) states, viewed as a cross-section from the side with full domains coloured, and from the top with only PLLs coloured. Orientation of black boxes (top) track orientation of core domains. **e.** Radius of the tunnel in open (green) and closed (magenta) states, measured using CHAP^54^. **f.** Phosphatidylethanolamine (PDB ID: 6OU) manually docked in the centre of the pore illustrates the size of a typical phospholipid compared to the pore diameter in the open and closed states. No density is observed for the lipid in our structures.

The configuration of the open and closed states is controlled primarily by two factors. First, the individual MCE domains from rings 1, 5, 6 and 7 undergo pivoting motions about axes through each MCE domain (**Fig. 4c**). Second, the PLLs of rings 1, 5, 6 and 7 shift, resulting in dilation and constriction of the corresponding pores, resembling the opening and closing of a camera diaphragm (**Fig. 4d**). The PLLs exhibit an additional layer of flexibility: within a given state (closed or open) the PLLs can likely adopt different conformations, as evidenced by additional loop-like densities in PLLs 5 and 7 (**Extended Data Fig. 7a**). Consistent with the observation that PLLs are flexible, a crystal structure we determined of the monomeric MCE2-MCE3 domains (residues 159-383) at 2.15 Å resolution (**Extended Data Table 2**) also revealed flexibility in the PLLs. The PLL is entirely disordered in MCE2 and adopts a significantly different conformation in MCE3 (**Extended Data Fig. 7b-f**), while the core MCE domains from the crystal structure are in good agreement with the EM structure. Our data suggest that while the tunnel may have a propensity to be in a more closed or more open conformation, which we were able to separate as discrete conformations, the PLLs are fluid in nature regardless of the overall conformation of the tunnel. These loop dynamics may facilitate the movement of molecules through the tunnel, and provide some plasticity to allow substrates of variable sizes to be accommodated. Interestingly, re-analysing our published cryo-EM dataset of MCE protein PqiB with newer versions of Relion revealed similar conformational changes in this smaller, three-ring barrel (**Supplementary Video 3, Extended Data Fig. 8a**). Together, the conformational changes result in overall modulation of the central pore diameter (**Fig. 4e, Extended Data Fig. 8a**) for both LetB and PqiB.

The opening and closing of the central pore through each MCE ring could play a role in tunnel gating and translocation of substrates. Phosphatidylethanolamine (PE) and phosphatidyl-glycerol (PG) constitute the bulk of the phospholipids in the OM^27,28^, and we have previously shown that LetB binds both PE and PG^7^. In the open state, the LetB tunnel is sufficiently wide to allow the passage of PE or PG from ring 1 to ring 6 (**Fig. 4e**). Docking these lipids into the pore of ring 6 reveals that in the open state these lipids are easily accommodated, while in the closed state their passage is restricted (**Fig. 4f**). The pore of ring 7 is narrower in both the closed and open states, and may restrict the flux of lipids to and from the OM. The density is weak for the tips of PLL7, where the tunnel is narrowest in ring 7, suggesting that these loops are very flexible and loop dynamics may facilitate lipid passage through the constriction.

### Pore-lining loops (PLLs) play a key role in LetB function

Since each PLL contributes to the formation of a dynamic, hydrophobic tunnel (**Fig. 2b**), we hypothesised that these loops may be essential for LetB function. To test this, we generated LetB PLL deletion mutants in which the β-hairpin motif of each PLL was deleted (**Fig. 5a**). To determine whether these mutants were functional in a cellular context, we again assessed whether these mutants could relieve the exacerbated phenotype of a *pqiAB letAB* double mutant in the presence of LSB. None of the LetB PLL mutants were functional in this assay, (**Fig. 5b**), suggesting that deleting any of the PLLs abolishes LetB function. Negative stain EM revealed that ΔPLL2 and ΔPLL3 form stable hexameric structures similar to the WT protein (**Fig. 5c**), while the other mutants appeared to be destabilised to some degree, as they assembled into hexameric structures at a lower frequency, or not at all (**Extended Data Fig. 6c**). Strikingly, 2D class averages of both ΔPLL2 and ΔPLL3 show a loss of density where the corresponding pore-lining loops were deleted (**Fig. 5c**), indicating that there is a hole in the wall of the tunnel.

**Figure 5.**
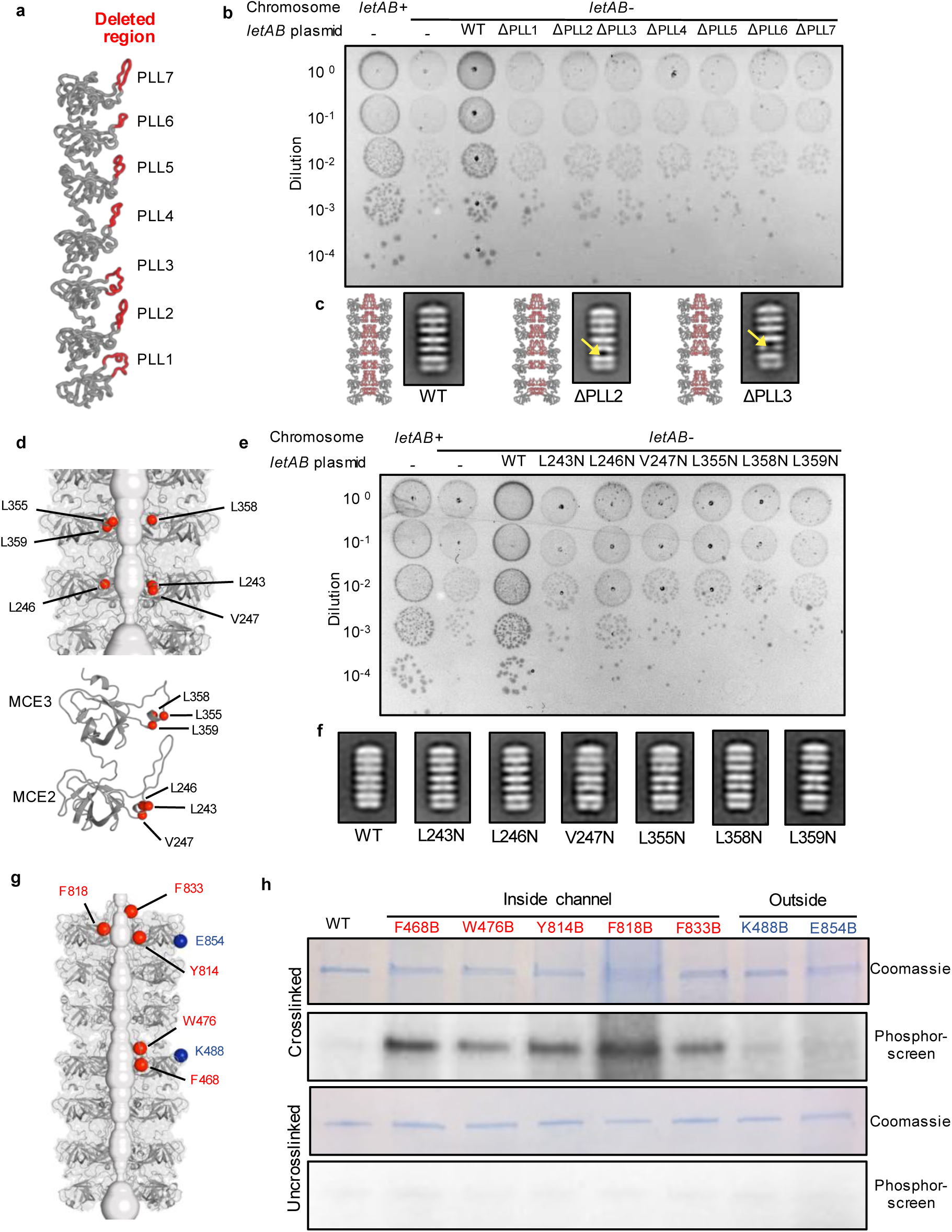
LetB tunnel is critical for substrate binding and function. **a-c.** Effect of PLL deletions on LetB structure and function. **a.** PLL regions deleted in LetB mutants (red) shown in the context of a LetB protomer. Seven mutants were made; in each the red region of a single PLL was deleted. **b.** Cellular assay for the function of LetB PLL deletion mutants. 10-fold serial dilutions of the indicated cultures spotted on plates containing LSB and incubated overnight. The *letAB* double mutant grows poorly in the presence of LSB, but can be rescued by the expression of *letAB* constructs containing WT *letB*. Expression of constructs with *letB* PLL deletion mutants fail to rescue. **c.** Negative stain 2D class averages of ΔPLL2 and ΔPLL3 mutants. Yellow arrows indicate missing density at site of deletion. A predicted structure of the hexamer is shown to the left of each 2D class aver age, in which the PLLs are in red and the deleted amino acids have been deleted from the model. **d-e.** Effect of point mutations in PLL2 and PLL3 on LetB structure and function. **d.** Mutated hydrophobic residues are shown as red spheres in the context of the tunnel (top) and zoomed in on each MCE domain (bottom). **e.** Cellular assay for the function of LetB with mutations in hydrophobic motifs. Results depicted as in part **b** above. Expression of constructs with indicated *letB* point mutations fail to rescue growth. **f.** Negative stain EM 2D class averages for each point mutant, showing similar structural organisation as the wild type. **g-h.** Substrates are crosslinked inside the LetB tunnel **g**. Residues in LetB that were mutated for incorporation of photo crosslinking amino acid, BPA are shown as spheres (red, inside the tunnel; or blue, outside the tunnel). **h**. SDS-PAGE analysis of purified LetB and BPA mutants, either crosslinked or uncrosslinked and stained by Coomassie (protein) or phosphor-imaged (^32^P signal).

Since our results show that ΔPLL2 and ΔPLL3 are folded yet functionally impaired, we asked whether specific amino acid residues within PLL2 and PLL3 may play specific roles in LetB function. An analysis of PLL sequences showed that each PLL contains a conserved hydrophobic motif, ΦxxΦΦ (Φ denotes a hydrophobic residue, × denotes any residue), which faces the tunnel interior. A similar hydrophobic motif is also observed in the MCE protein MlaD, and is important for function^7^. To test if these residues also play a functional role in LetB, we generated six point mutants: L243N, L246N, V247N in PLL2 and L355N, L358N, L359N in PLL3 (**Fig. 5d**). Our complementation assay showed that none of these mutants are able to relieve the exacerbated phenotype of a *pqiAB letAB* double mutant in the presence of LSB (**Fig. 5e**), suggesting that each point mutation in the hydrophobic motifs of PLL2 or PLL3 is sufficient to disrupt LetB function. Negative stain EM data of recombinantly produced mutants shows that each mutant is well-folded and assembled into a hexameric structure similar to the wild type (**Fig. 5f**), suggesting that the observed phenotypes likely correspond to a loss of function and not destabilisation of the proteins. Together, our results suggest that the intact hydrophobic tunnel lining formed by the PLLs is important for LetB function.

### Substrates are bound in the tunnel of LetB

Our data point to a model in which LetB transports substrates between the IM and OM through a central tunnel. Previous work has demonstrated that both LetB and its homologs are phospholipid-binding proteins^7,8,29,30^. However, it is unknown precisely where phospholipids or other substrates interact with LetB, and whether the interior of the tunnel is indeed part of the transport pathway. To assess whether potential substrates such as phospholipids are present in the tunnel, we developed a crosslinking-based approach to identify the locations of substrate-binding sites within LetB. We site-specifically incorporated the unnatural photocrosslinking amino acid p-benzoyl-L-phenylalanine (BPA) at multiple locations within LetB using amber stop codon suppression and an orthogonal aminoacyl-tRNA synthetase/tRNA system in *E. coli*^31^. UV irradiation of the sample leads to crosslinking of BPA to the C-H bonds of potential substrates nearby. To facilitate detection of these crosslinked adducts, E. coli cultures were grown in the presence of ^32^P orthophosphate, leading to labelling of bulk phospholipids and other phosphate-containing molecules. Thus crosslinking-dependent ^32^P association with LetB would be consistent with phospholipids being in close proximity to the site of BPA incorporation. As BPA is a bulky amino acid that is somewhat larger than tryptophan, we targeted all of the sites in the tunnel that presented bulky side chains facing the lumen for individual replacements with BPA (F468, W476, Y814, F818, F833) (**Fig. 5g**). In parallel, we designed mutations on the exterior, periplasm-facing surface of LetB where we hypothesised that phospholipids would not be present (K488, E854) (**Fig. 5g**).

After crosslinking in *E. coli* lysates, we purified the LetB protein by affinity chromatography, separated samples by SDS-PAGE, and detected LetB protein and associated phospholipids by Coomassie staining and a phosphorimager, respectively. Strong ^32^P association with LetB was detected at all five sites of BPA incorporation inside the tunnel (**Fig. 5h**), while minimal ^32^P association was detected at locations on the exterior surface of the tunnel. Thus, phospholipids or other ^32^P containing substrates are specifically present inside the LetB tunnel.

## Discussion

Our data support a model wherein LetB serves as a tunnel to mediate lipid transport (**Fig. 6**), providing a shielded environment for hydrophobic lipids to traverse the hydrophilic periplasm and the peptidoglycan (**Extended Data Fig. 8b**). LetB is sufficiently long to span the gap between the inner and outer membranes, and may physically interact with lipids and/or proteins in both membranes simultaneously. Consequently, the hydrophobic tunnel running the length of LetB may create a continuous pathway between the lipid environments of the IM and OM. To facilitate lipid insertion and extraction from membranes, it is likely that protein complexes in the IM and OM will also play a role. Indeed, the integral membrane protein LetA, which is encoded in the same operon as LetB, is a candidate to fulfill this role in the IM, while the existence of an OM complex remains hypothetical at this point.

**Figure 6.**
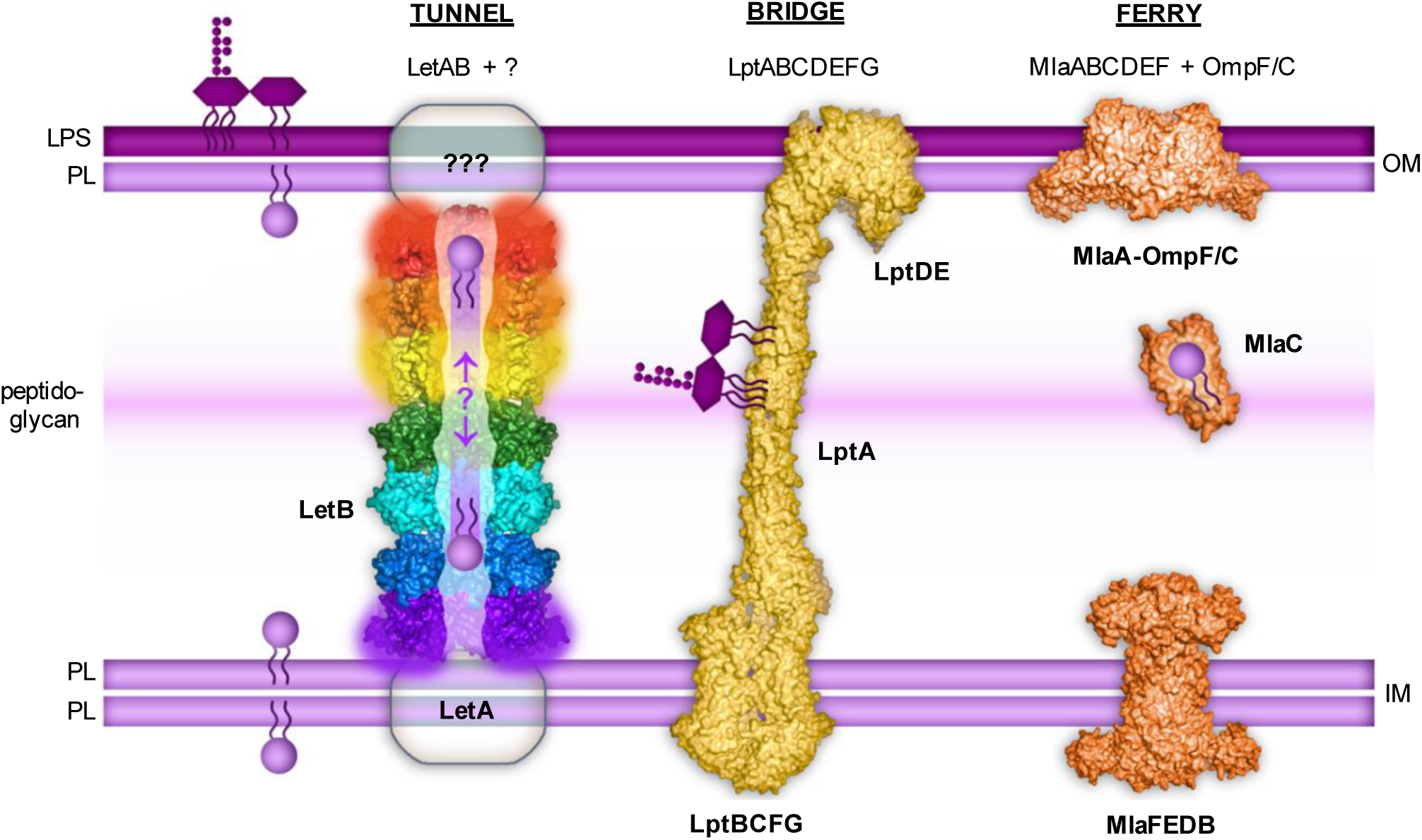
Model for LetB function in lipid transport. LetB forms a tunnel capable of spanning the periplasm, potentially interacting with LetA in the IM and unknown partners in the OM. Rings 1, 5, 6 and 7 are dynamic and adopt open and closed states (indicated by blurring). Lipids are transported through the tunnel, though the direction of transport remains unknown. Two other lipid transport systems in *E. coli* are depicted: 1) The LPS transport system, which is composed of Lpt proteins (PDB IDs: 5iv9, 2r1a and 6mit), forms a bridge between the two membranes along which LPS is transported; and 2) The Mla system, another MCE-protein family member (PDB IDs: 5nup, 5uwa and EMD 8610), which is composed of IM and OM complexes, and a soluble protein, MlaC, which “ferries” lipids across the hydrophilic periplasm. These three systems employ fundamentally different mechanisms for lipid transport, analogous to tunnels, bridges, and ferries.

This tripartite organisation (IM, periplasmic, and OM subunits) is a common feature of many transenvelope transport systems, where complexes in the IM and OM mediate the movement of substrates in, out, or across the membranes, while the periplasmic components facilitate transport in between. While in LetB, a single polypeptide is capable of spanning the periplasm, several other systems bridge the inner and outer membranes via protein-protein interactions of different subunits, and are evolutionarily unrelated to the MCE protein family. Among these are the Lpt system, which uses ATP hydrolysis to translocate the glycolipid LPS across an open, solvent exposed “bridge” of LptA proteins towards a protein complex in the OM^32–41^(**Fig. 6**). In contrast, other transenvelope transport systems employ a soluble periplasmic “ferry” protein to shuttle substrates between complexes in the IM and OM, as has been proposed for the lipoprotein export system Lol^32^, and a subset of MCE transporters, such as Mla^7^ (**Fig. 6**). Thus, while “bridges” of LptA are used for LPS transport, the MCE family of transporters appears to employ at least two distinct strategies - ferries and tunnels - for periplasmic transport of phospholipids.

In contrast to the Lpt system, whose function is well established as an exporter of LPS, the direction of transport mediated by MCE systems is less clear, as are the particular lipid species transported by the diverse MCE systems encoded in bacterial genomes. Whether these systems act on different substrates, under different environmental conditions, or perhaps even drive transport in different directions, remain open questions. While the membrane protein LetA may be predicted to fulfill a role similar to the ABC transporter permease MlaE, they appear to be evolutionarily unrelated, and may therefore function very differently from one another. Indeed, there is no evidence thus far that the Let system is driven by ATP hydrolysis, and we cannot rule out that other energy sources may be coupled to transport. Alternatively, the Let system may allow passive equilibration of lipids between the IM and OM, in a manner conceptually similar to the sites of IM-OM hemifusion that were proposed over 50 years ago^42^, but could potentially be mediated by transenvelope protein complexes instead of direct interaction of the IM and OM to facilitate lipid exchange. Our studies presented here provide insights into the architecture and mechanism of LetB, which forms a dynamic tunnel to transport lipids across the periplasm.

## Methods

### Phenotypic assays for MCE mutants in *E. coli*

All mutants were constructed in the background of *E. coli* K-12 BW25113. Bacteria were grown in LB medium (Difco) or on LB plates (Difco broth supplemented with 1.5% nutrient agar) and incubated at 37 °C. For strains harbouring a plasmid, the liquid medium was supplemented with 100 μg/ml carbenicillin. Strains with deletions of *pqiAB* and/or *letAB* were constructed in a previous study^9^. To test the function of various LetB mutants, the WT *letAB* operon from BW25113 was cloned into pET17b, and leaky uninduced expression of LetAB from this plasmid was sufficient to complement a *letAB* chromosomal deletion. LetB mutations of interest where introduced into this plasmid carrying both *letA* and *letB*, yielding a construct that expressed WT LetA and the desired LetB mutant. To test for functional complementation, 10-fold serial dilutions of overnight cultures of the relevant strains were prepared in a microtiter plate, and a multichannel pipette was used to spot 2 μl of each dilution to LB agar plates supplemented with sodium cholate (8%, Sigma Aldrich), sodium deoxycholate (4%, Sigma Aldrich), or lauryl sulfobetaine (LSB; (1%, Sigma Aldrich). Plates were incubated overnight at 37 C, then photographed using a GelDoc system (Bio-Rad). At least three independent colonies were used to perform replicates for each phenotypic assay.

### Expression and purification of LetB

DNA corresponding to the periplasmic portion of LetB (residues 43-877) was amplified from the MG1655 genome and cloned by Gibson assembly into a custom pET vector (pBEL587) resulting in a C-terminal 6xHis tag. The resulting plasmid (pBEL1324) was transformed into Rosetta 2 (DE3) cells (Novagen). For expression, overnight cultures of Rosetta 2 (DE3)/pBEL1324 were diluted 1:100 in LB (Difco) supplemented with carbenicillin (100 µg/mL) and chloramphenicol (38 µg/mL) and grown at 37 °C with shaking to an OD600 of 0.9, then induced by addition of IPTG to a final concentration of 1 mM and continued incubation overnight shaking at 15 °C. Cultures were harvested by centrifugation, and the pellets were resuspended in lysis buffer (50 mM Tris pH 8.0, 300 mM NaCl, 10 mM imidazole). Cells were lysed by two passes through an Emulsiflex-C3 cell disruptor (Avestin), then centrifuged at 38,000 xg to pellet cell debris. The clarified lysates were incubated with NiNTA resin (QIAGEN) at 4 °C, which was subsequently washed with Ni Wash Buffer (50 mM Tris pH 8.0, 300 mM NaCl, 10 mM imidazole) and bound proteins eluted with Ni Elution Buffer (50 mM Tris Ph 8.0, 300 mM NaCl, 250 mM imidazole). LetB containing fractions eluted from the NiNTA column were pooled and concentrated before separation on a Superdex 200 16/60 gel filtration column (GE Healthcare) equilibrated in 20 mM Tris pH 8.0 and 150 mM NaCl.

All mutant forms of the LetB periplasmic domain were produced by introducing the target mutation into pBEL1324 by site-directed mutagenesis and Gibson assembly of the resulting PCR products, and purified following similar procedures.

### Design of LetB truncation mutants

Two, three, four, five and six-ring truncation mutants of LetB were designed for both phenotypic assays and recombinant expression. For all mutant constructs, rings 1 and 7 were left intact, to preserve possible interactions with the inner and outer membranes/membrane proteins. Internal ring deletions were guided by cryo EM results (see **Results: Conformational dynamics modulate tunnel diameter)**, which showed that rings 2-4 are more rigid, likely functioning as one unit, while rings 5 and 6 are dynamic. For the six-ring truncation mutant we deleted ring 6; for the five-ring mutant we deleted rings 5 and 6; for the four-ring truncation mutant we deleted rings 2-4; for the three-ring truncation mutant we deleted rings 2-5. In the two-ring truncation mutant, all internal rings (rings 2-6) were deleted. For all expression constructs the TM helix of *letB* was removed. For phenotypic assays, full-length *letB* constructs were used.

### Negative stain electron microscopy of LetB

To prepare grids for negative stain EM, the sample was applied to a freshly glow discharged carbon coated 400 mesh copper grids and blotted off. Immediately after blotting, a 2% uranyl formate solution was applied for staining and blotted off. The stain was applied five times per sample. Samples were allowed to air dry before imaging. Data were collected on a Talos L120C TEM (FEI) equipped with a 4K × 4K OneView camera (Gatan) at a nominal magnification of 73,000x corresponding to a pixel size of 2 Å /pixel on the specimen, and a defocus range of 1 – 2 uM underfocus. Data processing was carried out in Relion 3.0^18^. Micrographs were imported, particles were picked manually followed by automated template-based picking. Several rounds of 2D classification were carried out using default parameters, except for the offset search range and offset search step, which were adjusted to 60 and 5 pixels, respectively. At least two replicates were performed for negative stain EM of each sample, starting from expression of recombinant protein.

### Cryo-EM grid preparation and data collection

After size exclusion chromatography on a Superdex 200 column (see Expression and Purification above), LetB was concentrated to 2 mg/mL. 3 μL of LetB at a final concentration of 0.5 mg/mL was applied to 400 mesh quantifoil holey carbon grids 1.2/1.3. The sample was then frozen in liquid ethane using the Vitrobot Mark III. Images were collected on a Titan Krios microscope (“Krios 2”) located at the HHMI Janelia Research Campus, operated at 300 kV and equipped with a Gatan K2 Summit direct electron detector camera. Images were recorded using SerialEM^43^ at a nominal magnification of 22,500x, corresponding to a pixel size of 1.31 Å. Further data collection parameters are shown in **Extended Data Table 1**.

### Cryo-EM data processing

Movies were drift corrected with MotionCor2^44^ and CTF estimation was performed using GCTF^45^. Data were processed with Relion 2.1^18,46^ unless otherwise stated, and the data processing workflow is described in **Extended Data Fig. 2**. ∼1,000 particles were selected manually and subjected to 2D classification. The resulting class averages were used as templates for subsequent automated particle picking. After 2D classification, 534,003 particles were selected from the 732,231 auto-picked particles and used for a first round of 3D classification. The map obtained from a reconstruction of the negatively stained sample^7^ (EMDB ID: EMD-8611) was used as a reference for initial 3D classification into three classes named cl_1 to cl_3 (left side of **Extended Data Fig. 2**). The class cl_1 with 226,082 particles led to a reasonable map with the expected size, which was selected for further processing. Maps from the other two classes were poorly defined, especially at the two ends of LetB, and were temporarily discarded. The best map was refined to ∼3.25 Å with a C6 symmetry and its local resolutions were estimated using *blocres* (kernel = 24 pixels) from Bsoft^47^. The following strategies were used to improve resolution. Most refinement was carried out with C6 symmetry imposed.

First, 3D classification into five classes of the whole map was performed. This resulted in classes dominated by different conformations in rings 5 to 7. In two of these classes, rings 5 to 7 adopt a closed conformation, and in two other classes, those rings adopt an open conformation where the tunnel diameter is larger. In the remaining class (cl_1.4 in **Extended Data Fig.2**), density for ring 1 was not visible and was weak even when the map was refined with no enforced symmetry.

Second, to improve the local resolution and analyse conformational heterogeneity, we masked, refined and classified different parts of the density map. Different sizes and types of masks were explored. The regions of interest were masked out and the projection of the high-resolution map subtracted from the micrographs. Iterative 3D classification and refinement was carried out as shown in **Extended Data Fig. 2**. The masking and signal subtraction of three consecutive rings was optimal, while refinement and classification of density corresponding to individual rings or of the pore-lining loops forming the central pore resulted in misalignment and poor resolution.

Finally, we investigated whether the resolution could be improved or new classes found by recovering the particles rejected by the initial 3D classification and using the maps obtained as starting references (right side of **Extended Data Fig. 2**). The whole process was repeated with the 534,003 particles originally selected after 2D classification instead of 226,082 and we thereby managed to further improve the resolution of the masked maps in the closed state.

The resolution of the best maps selected for *de novo* modelling varies from 2.96 Å to 3.78 Å. For each map, the overall resolution was estimated using the gold-standard Fourier Shell Correlation criterion (FSC=0.143). The FSC curves were calculated using soft masks and applying the masking-effect correction previously described^48^. The 3D FSC were computed using the “Remote 3DFSC Processing Server” at https://3dfsc.salk.edu/^49^. The local resolution maps were computed using Bsoft^47^.

To further analyse the relative movements of the MCE domains, we ran multi-body refinements in Relion 3.0-beta^25^ using model 1 (from map 1) of LetB. We tried non-overlapping masks of the 7 individual rings as well as masks of two consecutive rings with a one-ring overlap. Masking two consecutive rings with a one-ring overlap gave results that were interpretable. We also imported particles into CryoSparc v2.9^26^ to compute a 3D variability analysis using the map 1 obtained after 3D classification. Results are shown in **Supplementary Video 1**.

### Model building

The MCE domain of MlaD (PDB ID: 5UW2)^7^, was used to assign the hand of our LetB density map, and as a starting point for model building. The MlaD coordinates were initially fit into the density of the best resolved domain of LetB, (ring 3). The sequence was mutated to match that of ring 3 and residues were manually adjusted in COOT^50^. Then, the same process was repeated for all the other rings but using the model obtained for ring 3 as a reference. For each of the seven LetB rings, the initial model was built using the density map with the highest resolution (i.e. the masked map of the closed conformation for rings 1, 5, 6 and 7, and the wider masked density map for rings 2 to 4). The models were then subjected to the real_space_refine algorithm in PHENIX^51^ using 1 to 5 cycles and 50 to 100 iterations to optimise the fit and reduce clashes. These models were iteratively rebuilt in COOT then refined in PHENIX until completion. The models refined against these first maps were then fit into all the other maps and underwent rebuilding and refinement to model the observed alternative states of LetB.

Each MCE domain presents a PLL towards the lumen of the tunnel. As each loop roughly adopts a β-hairpin conformation, one would expect to observe two densities for each MCE domain (one for each strand of the β-hairpin), or 12 total densities for each hexameric MCE ring. However, for rings 2, 4, 7 and 5 in the closed conformation, 18 densities are visible in each ring, as it was observed for some of the rings of PqiB^7^. Due to the lower resolution of this part of the map, we were unable to identify whether the extra density corresponds to a substrate bound to the protein, or flexibility of the loops which may adopt multiple conformations. Since masking and 3D classification were unable to separate out different states or improve the maps, we built a single conformation of each loop through the densities that yielded the most sterically plausible final models.

We also generated “composite models” of LetB corresponding a fully closed (where the pores of the MCE1, 5, 6, and 7 rings are simultaneously all in their most constricted/closed state) and open state (where the pores of the MCE1, 5, 6, and 7 rings are simultaneously all in their most dilated/open state). As our classification and signal subtraction approaches generally divided LetB into three-ring modules, it is unclear if these fully open and fully closed states are well populated, or if some rings may adopt an open state at the same time others are closed. However, we find these models best illustrate the extent of the conformational changes in the transport tunnel. The composite closed model was created by combining the models built into map 5 (rings 1 and 2), 4 (rings 3 and 4), 3 (rings 5 to 7) and 1 (4 residues linking rings 4 to 5). The composite open model was created by combining the models built into map 8 (rings 1 and 2), 7 (rings 3 and 4), 6 (rings 5 to 7) and 2 (4 residues linking rings 4 to 5).

Statistics regarding the final models (**Table S2**) were extracted from the results of the real_space_refine algorithm as well as the the MolProbity^52^ and EMRINGER^53^ functions as implemented in the PHENIX package^51^. To display the movements between the closed and open configuration, we modified the ColorByRMSD plugin from pymol (pymolwiki.org/index.php/ColorByRMSD). To analyse the radii of the central pores, the CHAP software^54^. Molecular graphics and analyses were performed with either the UCSF Chimera package^55,56^ or the Pymol Molecular Graphics System (version 2.0 Schröodinger, LLC).

### Re-analysis of PqiB (EMDB ID: EMD-8612)

As many new cryo-EM software versions have recently become available with new capabilities, we recovered the data used previously to obtain the EM density map and model of PqiB (PDB 5UVN)^7^ and reanalysed it in Relion 2.1 Iterative 3D classification with masking and signal subtraction led to the identification of distinct closed and open states. When the classification is focused on the three MCE domains, we distinguish two clearly different classes: class 1 with 40,462 particles, and class 2 with 15,811 particles. These classes were refined at 3.8 Å and 4.4 Å respectively in which we fitted the model 5UVN. After a rigid-body docking of each MCE domain, the pore-lining loops of rings 1 and 2 of class 2 were further refined in Phenix. Multi-body refinement was carried out as described above for LetB.

### Expression and purification of LetB fragment for X-ray crystallography

DNA corresponding to the MCE2+MCE3 domains of LetB (residues 159-383) was amplified from the MG1655 genome and cloned by Gibson assembly into a custom pET vector (pDCE587) resulting in a C-terminal 6xHis tag. The resulting plasmid (pBEL1584) was transformed into Rosetta 2 (DE3) cells (Novagen). For expression, overnight cultures of Rosetta 2 (DE3)/pBEL1584 were diluted 1:100 in LB (Difco) supplemented with carbenicillin (100 µg/mL) and chloramphenicol (38 µg/mL) and incubated at 37 °C with shaking at 225 RPM to an OD600 of 0.9, then induced by addition of IPTG to a final concentration of 1 mM with continued shaking at 37 °C for 4 hours. Cultures were harvested by centrifugation, and the pellets were resuspended in lysis buffer (20 mM Tris pH 8.0, 300 mM NaCl). Cells were lysed by two passes through an Emulsiflex-C3 cell disruptor (Avestin), then centrifuged at 38,000 xg to pellet cell debris. The clarified lysates loaded onto a cOmplete His-tag Purification column (Roche) at 4 °C on an Akta FPLC (GE Healthcare), which was subsequently washed/eluted with a gradient of increasing concentrations of imidazole. LetB containing fractions eluted from the NiNTA column were pooled and concentrated before separation on a Superdex 200 16/60 gel filtration column (GE Healthcare) equilibrated in 20 mM Tris pH 8.0 and 150 mM NaCl.

### Crystal structure LetB MCE2-MCE3 domains

Size exclusion fractions containing LetB (residues 159-383, plus C-terminal His tag) were concentrated to 23 and 70 mg/mL, and sitting drop, vapor-diffusion crystallisation trials were conducted using the JCSG Core I-IV screens (QIAGEN). Crystals grew from drops consisting of 100 nL protein plus 100 nL of a reservoir solution consisting of 0.1 M Tris pH 8.5, 20% w/v PEG 1K, and were cryoprotected by supplementing the reservoir solution with 15% ethylene glycol. Native diffraction data was collected at the GM-CA/CAT beamline 23-ID-B at the Advanced Photon Source, and indexed to P6_5_ and reduced using XDS (**Extended Data Table 2**) (Kabsch, 2010). This dataset was phased by molecular replacement using Phaser (McCoy et al., 2007), using the MCE2 and MCE3 domains from the cryo-EM structure as search models. The resulting model was adjusted in Coot^50^ and refined using Phenix^51^. This final model consists of two copies of the MCE2-MCE3 domain fragment of LetB in a monomeric state.

### Crosslinking lipids in the tunnel of LetB

T7express *E. coli* (NEB) were co-transformed with plasmids to express periplasmic LetB (either WT or mutant forms derived from pBEL1324) and pEVOL-pBpF (Addgene #31190), which encodes a tRNA synthetase/tRNA pair for the *in vivo* incorporation p-benzoyl-l-phenylalanine (BPA) in *E. coli* proteins at the amber stop codon, TAG ^9,31^. Bacterial colonies were inoculated in LB broth supplemented with carbenicillin (100 µg/mL) and chloramphenicol (38 µg/mL) and grown overnight at 37 °C. The following day, bacteria were pelleted and resuspended in ^32^P Labelling Medium (a low phosphate minimal media we optimised starting from LS-5052^57^: 1 mM Na2HP04, 1 mM KH2PO4, 50 mM NH4Cl, 5 mM Na2SO4, 2 mM MgSO4, 20 mM Na2-Succinate, 0.2x trace metals and 0.2% glucose) supplemented with carbenicillin (100 µg/mL) and chloramphenicol (38 µg/mL) and inoculated 1:50 in the 10 ml of the same medium. Bacteria were grown to OD 0.9 and a final concentration of 1 mM IPTG, 0.2% L-arabinose and 0.5 mM BPA (Bachem), alongside 500 µCi ^32^P orthophosphoric acid (PerkinElmer) and left to induce overnight. The following day, the cultures were spun down and resuspended in 1 ml of freeze-thaw lysis buffer (50 mM Tris pH 8.0, 300 mM NaCl, 10 mM imidazole, 1 mg/ml lysozyme, 0.5 mM EDTA, 25U benzonase) and then underwent eight freeze-thaw cycles by alternating between liquid nitrogen and a 37 °C heat block. Unbroken cells were pelleted at 12,000 xg for 2 mins and the lysates were transferred to a 12 well culture plate. Samples to undergo crosslinking were treated with 365 nM UV in a Spectrolinker for 30 mins. Each sample was then mixed with 25 µl of nickel beads (Ni Sepharose 6 Fast Flow) for 30 mins. The beads were pelleted at 500 xg for 1 min and the supernatant collected. The beads were then washed four times with detergent wash buffer (50 mM Tris pH 8.0, 300 mM NaCl, 40 mM imidazole, 0.25% triton X-100) and finally resuspended in 50 µl of 2x SDS-PAGE loading buffer. The beads were boiled for 5 mins to elute protein and spun down at 12,000 xg for 2 mins. Eluted protein was analysed by SDS-PAGE and stained using InstantBlue™ Protein Stain (Expedeon). Relative LetB loading on the gel was estimated integrating the density of the corresponding bands in the InstantBlue-stained gel in ImageJ^58^, and this was used to normalise the amount of LetB loaded on a second gel, to enable more quantitative comparisons between samples. The normalised gel was stained and ^32^P signal was detected using a phosphor screen and scanned on a Typhoon scanner (Amersham). Three replicates of the experiment were done, starting with protein expression.

## Data availability

All cryo-EM and X-ray models and maps are temporarily available at our website: http://bhabhaekiertlab.org/pdb-links, and maps are available on request. All cryo-EM density maps of *E. coli* LetB will be deposited in the Electron Microscopy Data Bank, and the corresponding coordinates will be deposited in the PDB. The structure factors and coordinates for the crystal structure of LetB MCE2-MCE3 will be deposited in the PDB. Cryo-EM data for LetB will be deposited in EMPIAR. Plasmids will be deposited in Addgene.

## Acknowledgements

We would like to thank Kristen Dancel-Manning for help with preparing Figure 6 and EM training, James Fraser, Timothy Knowles, Iris Grossman-Haham, Juliana Ilmain, Stefan Niekamp and Rachel Redler for critical reading and feedback on our manuscript, and members of the Bhabha/Ekiert labs for helpful discussions. We gratefully acknowledge the following funding sources: R00GM112982 (G.B.), DFS-20-16 (G.B.), R35GM128777-01 (D.E), Summer Undergraduate Research Program at the NYU School of Medicine (CTM). We thank Chuan Hong and Zhiheng Yu at the HHMI Janelia Research Campus for assistance in microscope operation and data collection of cryo samples. GM/CA@APS has been funded in whole or in part with Federal funds from the National Cancer Institute (ACB-12002) and the National Institute of General Medical Sciences (AGM-12006). This research used resources of the Advanced Photon Source, a U.S. Department of Energy (DOE) Office of Science User Facility operated for the DOE Office of Science by Argonne National Laboratory under Contract No. DE-AC02-06CH11357. The Eiger 16M detector was funded by an NIH–Office of Research Infrastructure Programs, High-End Instrumentation Grant (1S10OD012289-01A1). We thank Fengxia Lang for overseeing use of the TalosL120C microscope; the Microscopy Core in which the TalosL120C is housed is partially supported by NYU Cancer Center Support Grant NIH/NCI P30CA016087. We thank the Central Lab Services team at the Skirball Institute supervised by Michael Mirabile for preparation of media and buffers in large quantities. EM data processing has utilised computing resources at the HPC Facility at NYU, and we thank the HPC team including Martin Ossowski, Ali Siavosh-Haghighi, Michael Costantino, Paul Glick, Loren Koenig, Nam Pho as well as Joe Katsnelson and John Speakman for high performance computing support.

## Author information

### Contributions

GB, DE, GI and NC conceived the research. All authors performed experiments, collected data and analysed data. GI, NC, GB and DE wrote the manuscript. All authors edited the manuscript.

### Materials and Correspondence

Correspondence and material requests should be addressed to Gira Bhabha or Damian C. Ekiert.

