## Extended and Supplementary info for "Structure of LetB reveals a tunnel for lipid transport across the bacterial envelope"

#### Extended Data Figure 1

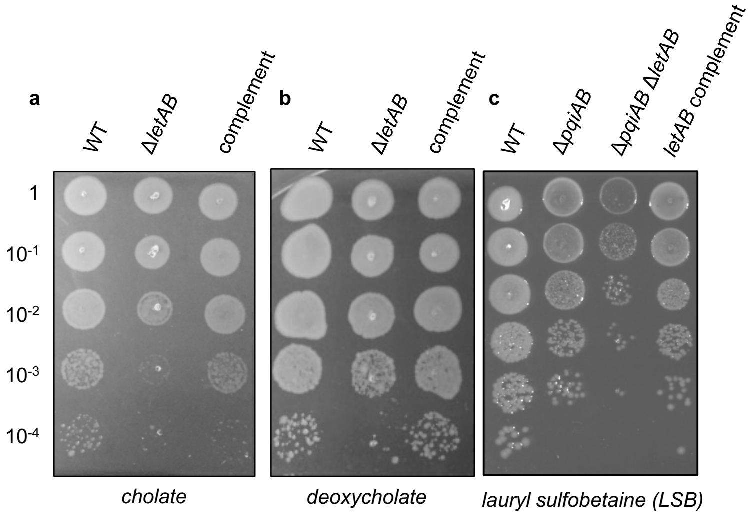


**Extended Data Figure 1. Phenotypes of *E. coli letAB* mutants show outer membrane defect.** Cellular assay for the function of LetB. 10-fold serial dilutions of the indicated cultures spotted on plates containing cholate (**a**) or deoxycholate (**b**) and incubated overnight. The *letAB* deletion mutant grows poorly in the presence of cholate (**a**) and deoxycholate (**b**), and can be rescued by complementation with a plasmid carrying WT *letAB*. **c**. The *pqiAB letAB* double mutant plated in the presence of LSB and grown overnight. *letAB* deletion exacerbates the *pqiAB* phenotype, which can be complemented with a plasmid carrying WT *letAB*. The growth defect on LSB is the most robust of the *letAB* phenotypes, and hence is used as our primary assay for LetB function.

### Extended Data Figure 2

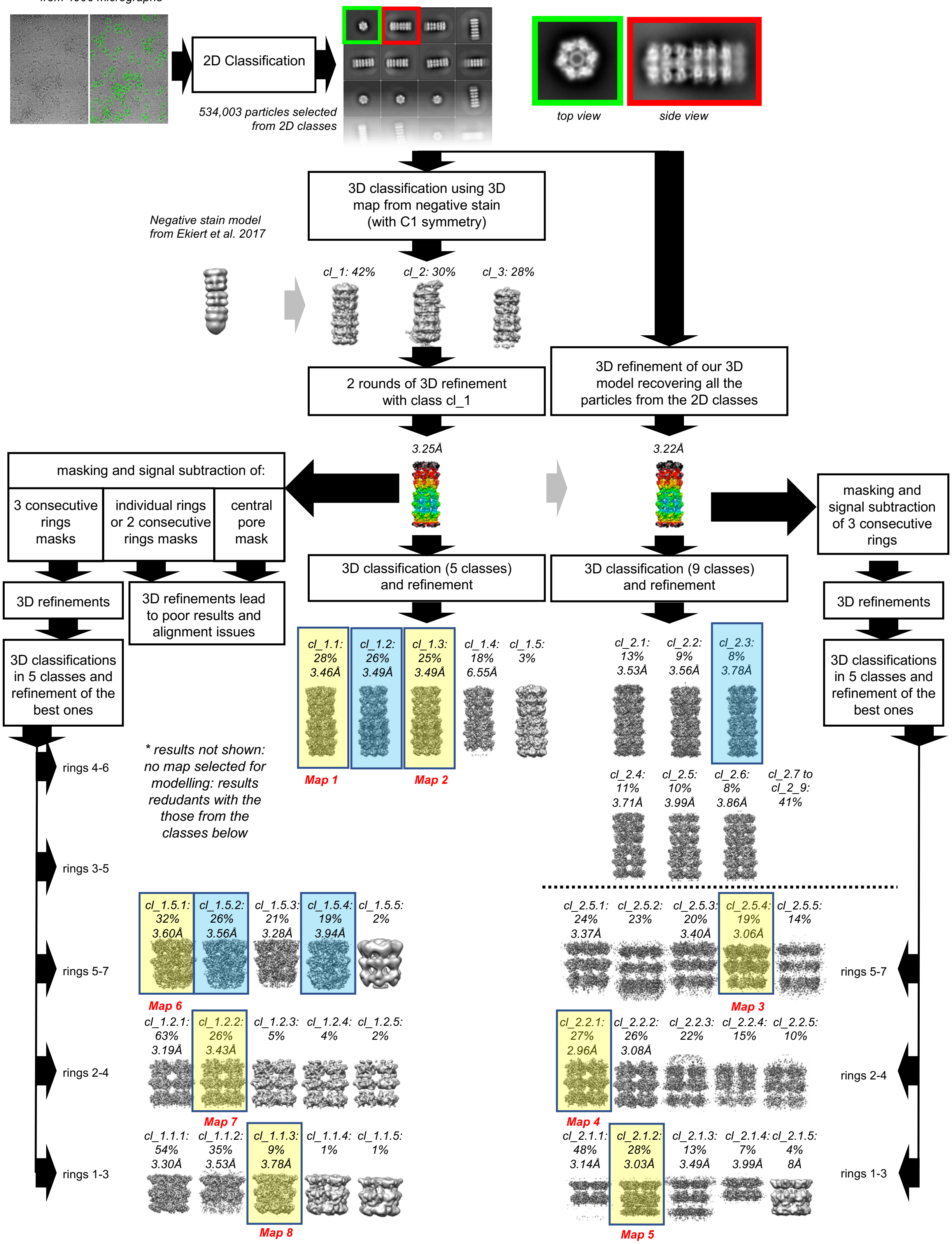


**Extended Data Figure 2. Cryo-EM data processing workflow.** Overall scheme for 3D classification, signal subtraction, masking and refinement. Yellow boxes indicate the maps used for model building and conformational dynamics analysis of LetB, and the blue boxes show other high resolution classes with minor differences in the open and closed states, not used for analysis. See **Methods** for more details.

### Extended Data Figure 3

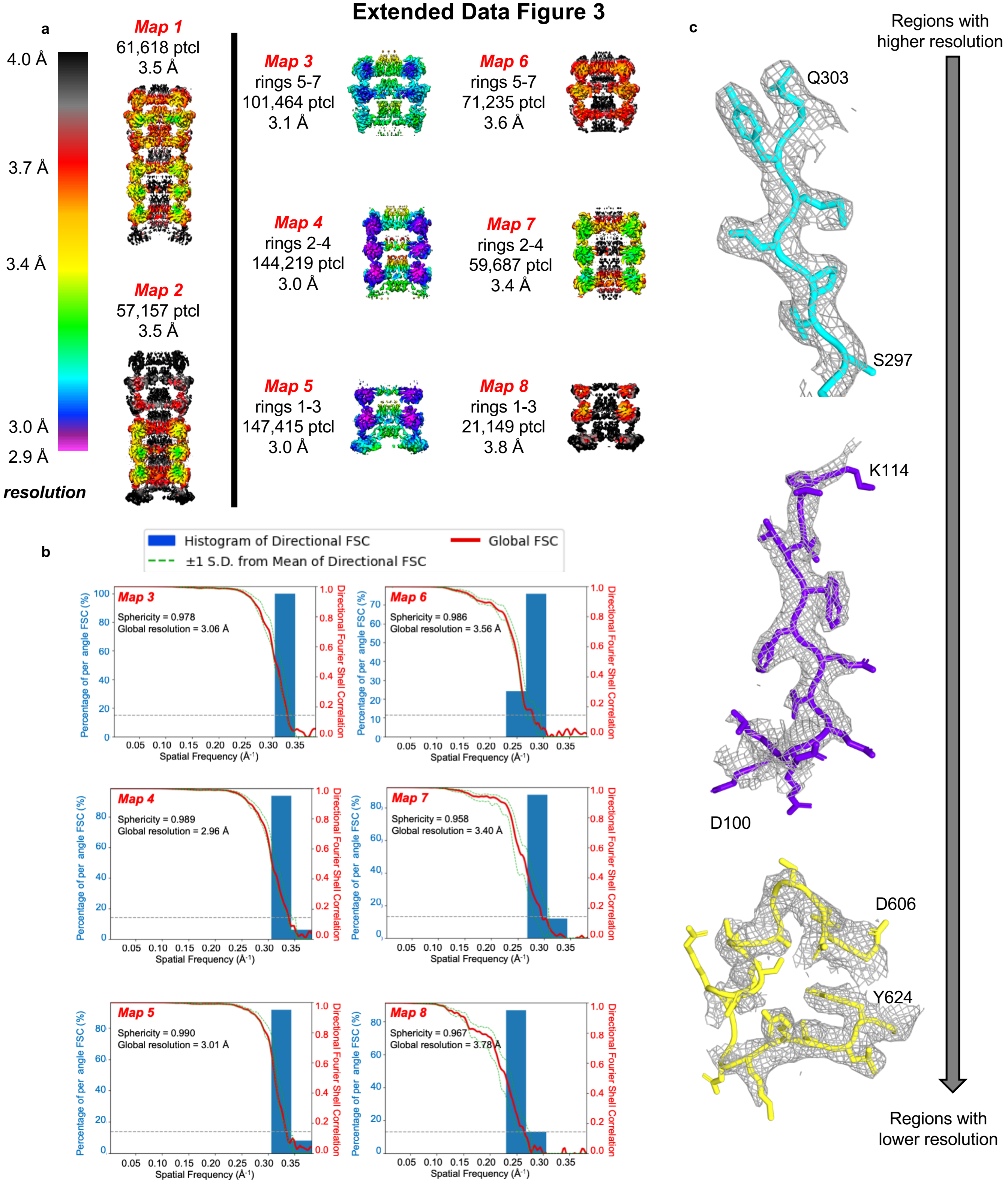


**Extended Data Figure 3. Overview of cryo-EM density map quality. a**. Cross-sectional views of density maps of LetB coloured by local resolution, as estimated using the blocres program from Bsoft[47]. Classes obtained prior to signal subtraction are shown on the left and improved maps of selected regions after signal subtraction are shown on the right. **b**. Fourier Shell Coefficient (FSC) and 3D FSC curves measured by the Gold-standard method (using the 3DFSC processing server[49]). **c**. Examples of density into which the model was built. Representative density in higher, intermediate and lower resolution regions are shown.

### Extended Data Figure 4

**a**

```
                     β1                      β2          β3
               1          11         21         31         41         51
RMSD: ca
LetB, MCE 1  46 .GNTVTIDFM .SA.DGIV.P GRTPVRYQGV EVGTVQD.IS L........S ..D.D.LRKI  88
LetB, MCE 2 160 .DLMIHLQAP .DL.GSLN.S G.SLVYFRKI PVGKVYD.YA I........N PNK....QGV 201
LetB, MCE 3 279 RGVIIKLELP .SG.AGLT.A DSTPLMYQGL EVGQLTK.LD L........N P.G..G..KV 321
LetB, MCE 4 391 DVLTLTLTAP .ESYG.IDAG ..QPLILHGV QVGQVID.RK L........T ..S.K...GV 431
LetB, MCE 5 515 ..TTVSLSAE .T.LP.DVQA G.SVVLYRKF EVGEVIT.VR P........R .....A.NAF 553
LetB, MCE 6 634 .GGQITLHAF .DA.GKLA.V G.MPIRYLGI DIGQIQT.LD L........I TA..R..NEV 675
LetB, MCE 7 746 DGLSIIVEAP .EA.GSLG.I G.TPVLFRGL EVGTVTG.MT L........G TLS....DRV 788
PqiB, MCE 1  42 .GPEVTLITA .NA.EGIE.G GKTTIKSRSV DVGVVES.AT L........A DDL....THV  84
PqiB, MCE 2 156 KGIRVILDSK .KA.GQLS.P G.DPVLFRGY RVGSVET.ST F........D TQ..K..RNI 198
PqiB, MCE 3 284 DHIDYLMFFK DSV.RG.LQP G.APVEFRGI RLGTVSKVPF .FAPNMRQTF .N.DD..YRI 335
MlaD         38 PTYTLYATFD .N.IGGLK.A R.SPVSIGGV VVGRVAD.IT L........D PK..T..YLP  80

               β4         H4                   β5          PLL
               61         71         81         91         101        111
RMSD: ca
LetB, MCE 1  89 EVKVSIKSD. MK..D..A.L R.EE...TQF WLVTP.KASL AGVSG..... ...LDALVGG 129
LetB, MCE 2 202 VIDVLIERR. FT..D..L.V K.KG...SRF WNVSG.VDAN VSISG.AKVK LESLAALVN. 248
LetB, MCE 3 322 TGEMTVDPS. VV..T..L.L R.EN...TRI ELRNP.KLSL SDAN...... ...LSALLTG 361
LetB, MCE 4 432 TFTVAIEPQ. HR..E..LVK G..D...SKF VVNSR.VDVK VGLDGVEFLG AS.ASEWIN. 478
LetB, MCE 5 554 DIDLHIKPE. YR..N..LL. T.SN...SVF WAEGGAKVQL NGSGLTVQAS P..LSRALK. 600
LetB, MCE 6 676 QAKAVLYPE. YV..Q..TFA R.GG...TRF SVVT..PQIS AAGVEH.... ...LDTILQ. 716
LetB, MCE 7 789 MIAMRISKR. YQ..H..L.V R.NN...SVF WLASG.YSLD FGLTGGVVKT GT.FNQFIR. 835
PqiB, MCE 1  85 EIKARLNSG. ME..K...LL H.KD...TVF WVVKP.QIGR EGISG..... ...LGTLLSG 125
PqiB, MCE 2 199 SYQLFINAP. YD..R..L.V T.NN...VRF WKDSG.IAVD LTSAGMRVEM GS.LTTLLS. 245
PqiB, MCE 3 336 PVLIRIEPER L.KDVVEH.L GELLKRGLRG SLK....... .......... ...TGNLVTG 373
MlaD         81 RVTLEIEQR. Y...N..HI. P.DT...SSL SIR....... .......... ...TSGLLG. 108

                    β6                                        β7
               121        131        141        151        161
RMSD: ca
LetB, MCE 1 130 .NYIGMMPG. .......K.. GK.EQ....D .......... HFV.ALD.   149
LetB, MCE 2 249 .GAIAFDSP. ..E...ES.. .KPAEA..ED .......... TFG.LYE.   272
LetB, MCE 3 362 .KTFELVPG. .......D.. GE.PR....K .......... EFV.VVPG   382
LetB, MCE 4 479 .GGIRILPG. ......DK.. GE.MK....A .......... SYP.LYA.   499
LetB, MCE 5 601 .GAISFD.NL ..S....... .......... GASASQRKGD KRI.LYA.   625
LetB, MCE 6 717 .PYINVEPG. .......R.. GN.PR....R .......... DFE.LQEA   737
LetB, MCE 7 836 .GGIAFA.TP ..PGTPL... APKAQE..GK .......... HFL.LQE.   862
PqiB, MCE 1 126 .VYIELQPG. ......AK.. GS.KM....D .......... KYD.LLDS   147
PqiB, MCE 2 246 .GGVSFD.VP EGLD..LG.. .QPVAP..KT .......... AFV.LYDD   273
PqiB, MCE 3 374 ALYVDLD... .......... ......NGYQ .......... IIP.TVSG   391
MlaD        109 EQYLALNVGF ......E.LG TAILKD..GD .......... TIQDTKS.   136
```

**b**

% identity

| protein | domain | residues | LetB MC1 | LetB MCE2 | LetB MCE3 | LetB MCE4 | LetB MCE5 | LetB MCE6 | LetB MCE7 | PqiB MCE1 | PqiB MCE2 | PqiB MCE3 | MlaD MCE1 |
|---|---|---|---|---|---|---|---|---|---|---|---|---|---|
| LetB | MCE1 | 46-149 | 100.0 | 19.4 | 29.7 | 17.4 | 20.0 | 28.4 | 24.3 | 37.5 | 23.3 | 18.9 | 21.4 |
| | MCE2 | 158-272 | | 100.0 | 14.6 | 28.4 | 28.8 | 15.4 | 32.2 | 19.1 | 29.6 | 17.5 | 17.4 |
| | MCE3 | 279-382 | | | 100.0 | 16.5 | 15.2 | 29.7 | 24.3 | 19.4 | 19.4 | 12.1 | 14.8 |
| | MCE4 | 391-499 | | | | 100.0 | 21.9 | 17.4 | 26.6 | 18.2 | 18.4 | 13.2 | 24.2 |
| | MCE5 | 513-625 | | | | | 100.0 | 16.0 | 24.6 | 20.8 | 29.0 | 15.8 | 17.7 |
| | MCE6 | 634-737 | | | | | | 100.0 | 20.2 | 26.9 | 18.3 | 11.1 | 11.4 |
| | MCE7 | 746-862 | | | | | | | 100.0 | 21.9 | 33.1 | 21.2 | 20.4 |
| PqiB | MCE1 | 42-147 | | | | | | | | 100.0 | 21.9 | 12.1 | 13.5 |
| | MCE2 | 156-273 | | | | | | | | | 100.0 | 23.2 | 21.4 |
| | MCE3 | 284-412 | | | | | | | | | | 100.0 | 18.9 |
| MlaD | MCE1 | 38-139 | | | | | | | | | | | 100.0 |
| | | min | 17.4 | 14.6 | 12.1 | 13.2 | 15.2 | 11.1 | 20.2 | 12.1 | 18.3 | 11.1 | 11.4 |
| | | avg | 24.0 | 22.2 | 19.6 | 20.2 | 21.0 | 19.5 | 24.9 | 21.1 | 23.7 | 16.4 | 18.1 |
| | | max | 37.5 | 32.2 | 29.7 | 28.4 | 29.0 | 29.7 | 33.1 | 37.5 | 33.1 | 23.2 | 24.2 |

**Extended Data Figure 4. Sequence comparison of MCE domains from *E. coli*. a**. Alignment of the *E. coli* MCE domains from MlaD, PqiB and LetB proteins, generated using ClustalO[60] from JalView 2[61]. **b**. Pairwise percent identity between all MCE domains, coloured from red (lowest) to blue (highest).

#### Extended Data Figure 5

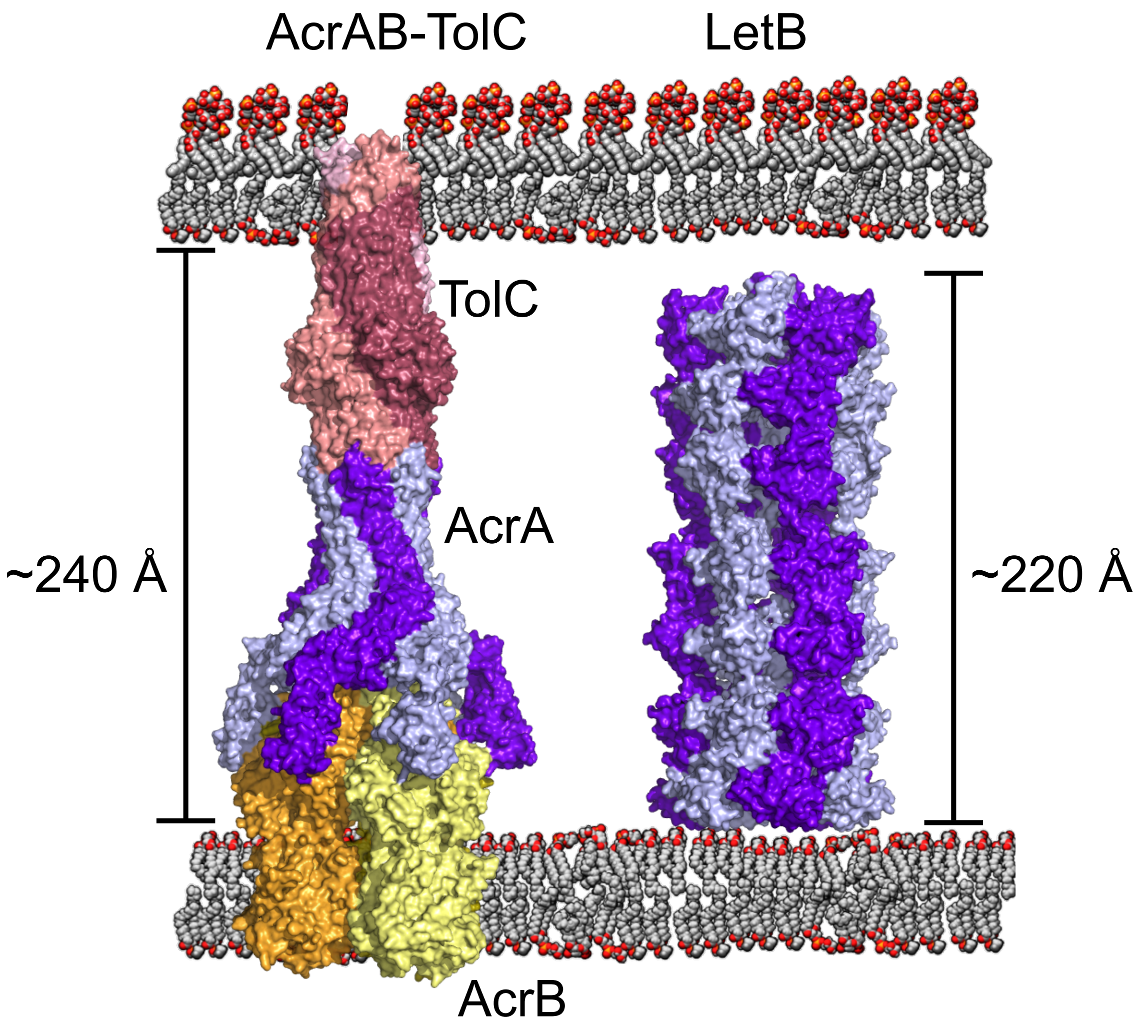


**Extended Data Figure 5. Comparison of LetB with AcrAB-TolC.** Surface representations of AcrAB-TolC (PDB ID: 5o66) and LetB, coloured by protein subunit. The periplasmic regions of LetB and AcrAB-TolC are of similar lengths, ~220 and ~240 Å respectively, consistent with the length of the periplasm. The periplasmic width (240 Å) is shown based on the hydrophobic regions of AcrAB-TolC[59], and is close to other reported values of 210[22] and 230 Å[21]. The periplasmic space is known to vary, for example in response to stress[62], and may not be constant in all regions of the cell envelope.

**Extended Data Figure 6**

a

| No. MCE | Gene name | Species |
|---|---|---|
| **4xMCE** | UWK_02604 | *Desulfocapsa sulfexigens* |
| **5xMCE** | D791_00999 | *Nitrincola nitratireducens* |
| **6xMCE** | PA4689 | *Pseudomonas aeruginosa* |
| **7xMCE** | *letB (yebT)* | *Escherichia coli* |
| **8xMCE** | Dacet_0211 | *Denitrovibrio acetiphilus* |

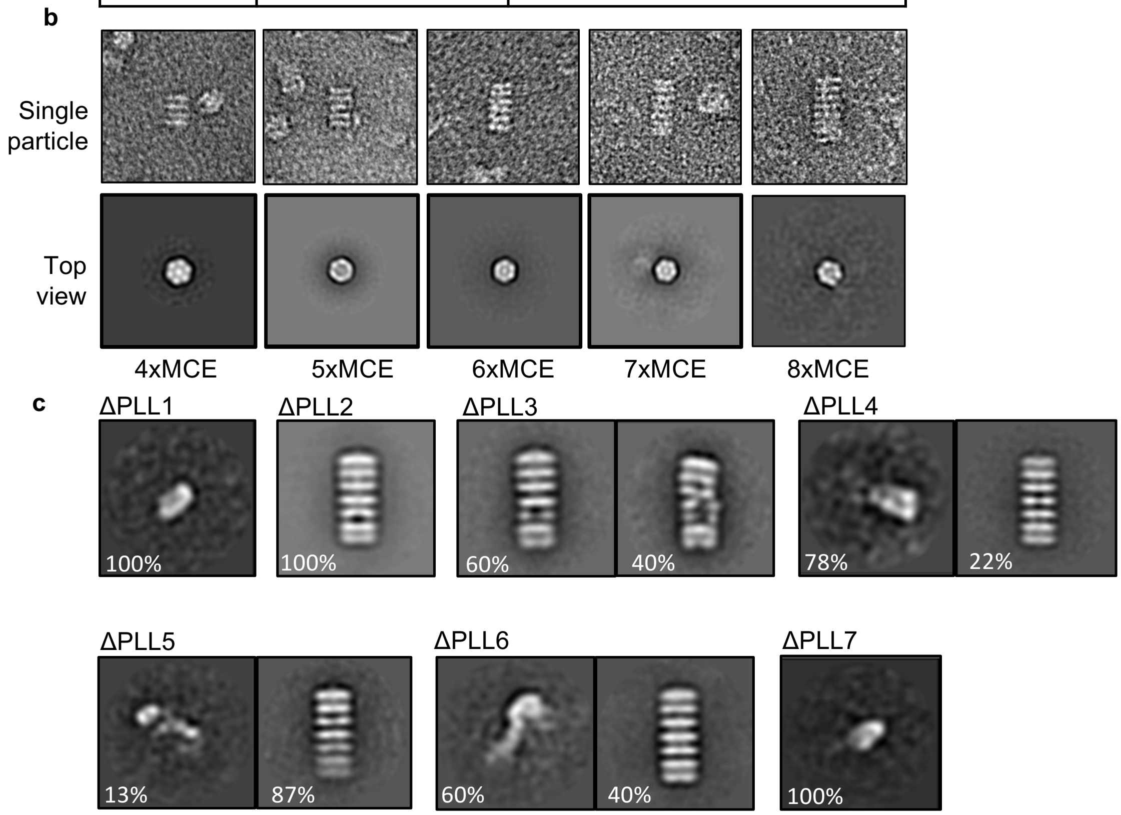


**Extended Data Figure 6. Negative stain EM data for naturally occurring proteins with varying number of MCE domains and LetB PLL mutants. a**. Gene IDs of naturally occurring MCE proteins of varying size and the corresponding bacterial species of origin. **b**. Single particles and top views of 2D class averages of naturally occurring proteins with varying number of MCE domains. **c**. 2D class averages of PLL-deletion mutants, related to **Fig. 5 a-c**. For completeness, classes shown for ΔPLL2 and 60% particles for ΔPLL3 are duplicated from **Fig. 5c**.

### Extended Data Figure 7

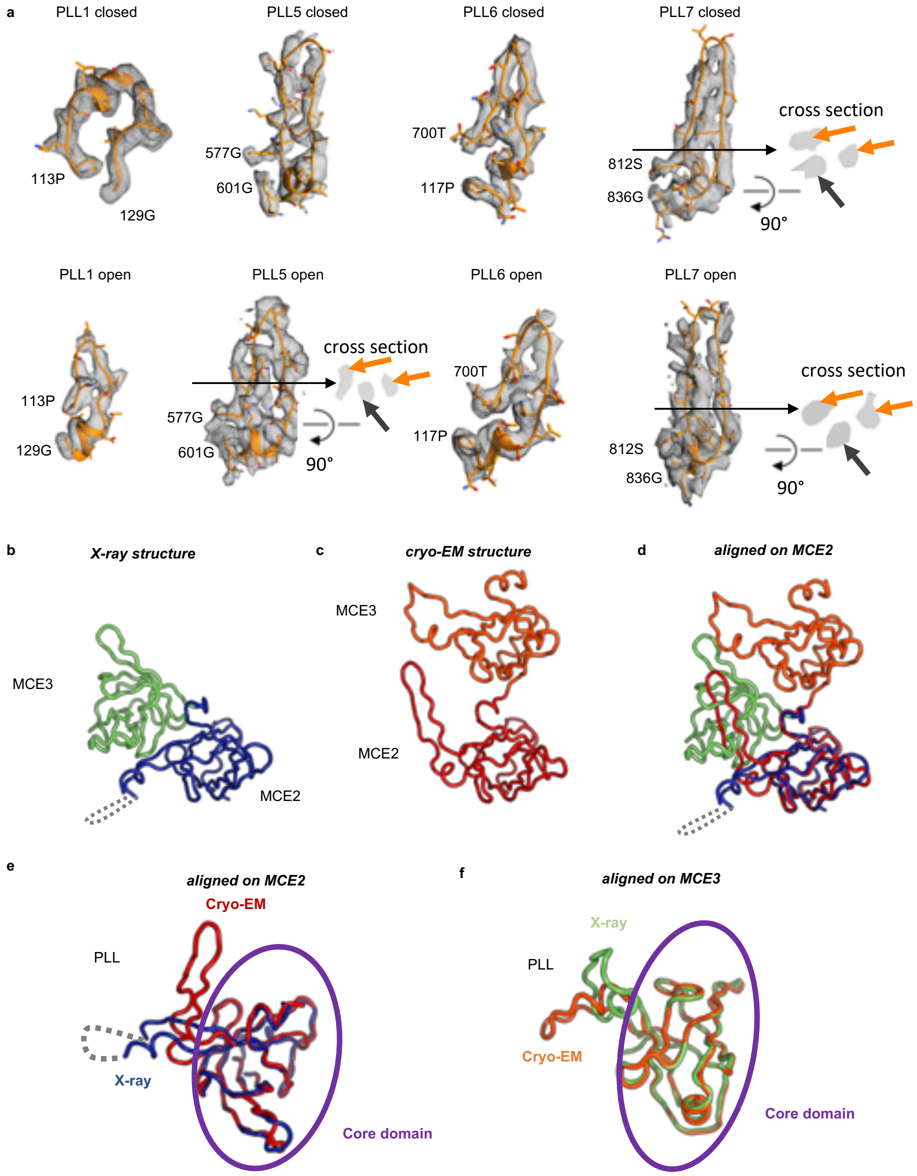


**Extended Data Figure 7. Multiple conformations of pore-lining loops (PLLs)**. **a**. Densities for PLLs of rings 1, 5, 6 and 7, which change conformations between the closed and open states. An extra “third” density is observed in the region of PLL5 in the open state and PLL7 in both states, indicated by a black arrow in the cross sections. This likely represents an alternate conformation of the loop, but due to the low resolution, we have not modeled additional states explicitly. Additional density is similarly observed in the regions of PLL2 and PLL4 (not shown). **b.** X-ray structure of LetB MCE2-MCE3 monomer fragment. **c**. Cryo-EM structure showing only LetB MCE2-MCE3 of one monomer. **d-f**. Comparison of MCE2-MCE3 domains from X-ray and cryo-EM structures. **d,e**. Superposition on the MCE2 domain, showing large-scale domain movement of MCE3 (**d**) and differences in PLL2 conformation (**e**) in the monomeric X-ray structure compared with the hexameric cryo-EM structure. The relative orientations of the MCE2 and MCE3 domains (**d**) are quite different from the state observed in the cryo-EM structure, likely due to the difference in oligomeric state and the flexible linker/interface between the two domains. Superposition of core domains show excellent agreement between the X-ray and cryo EM structures in these regions (RMSD: 0.6 for MCE2 core domain and 0.7 for MCE3 core domain). In addition, the partial disorder of PLL2 in the X-ray structure (**e**) suggests that the PLLs are inherently flexible. **f.** Superposition on the MCE3 domain showing that PLL3 is in different conformations in the X-ray and cryo-EM structures.

#### Extended Data Figure 8

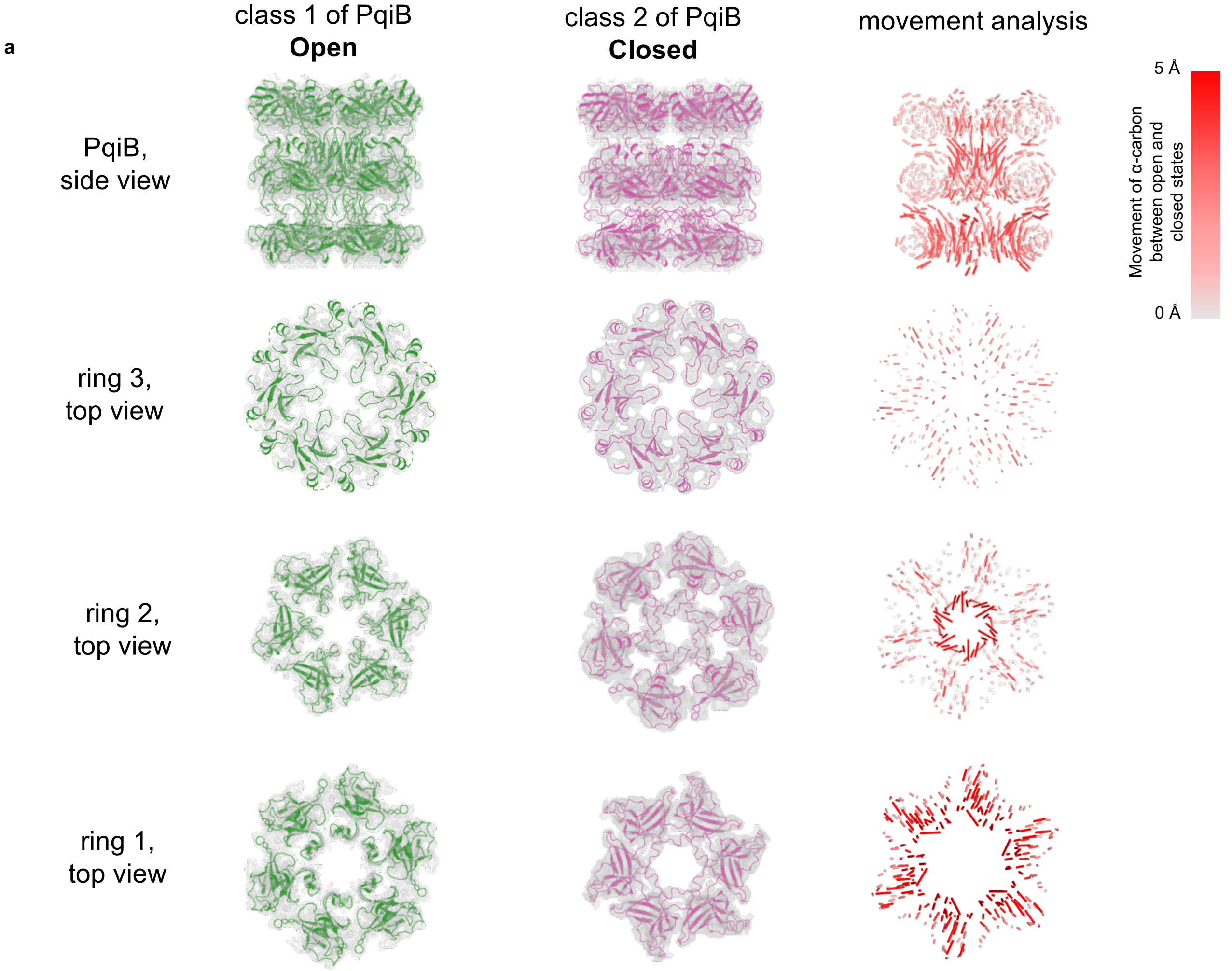


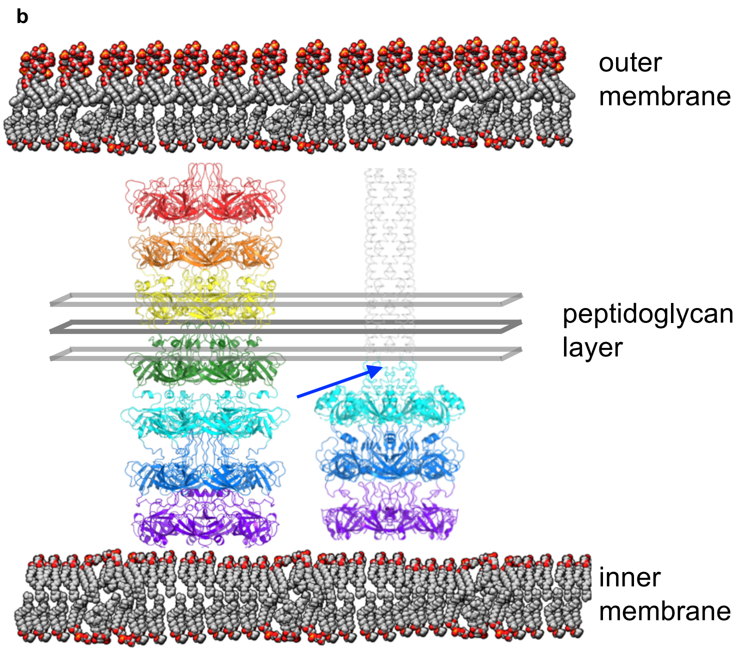


**Extended Data Figure 8. Conformational dynamics in PqiB. a**. Re-analysis of previously published PqiB data[7] shows open and closed states similar to those observed for LetB. Side and top views are shown for the open (green) and closed (magenta) states, with maps represented as grey mesh. Distances between Cα atoms in each state is mapped in the movement analysis, with longer lines and red colour representing regions of greater displacement. **b**. LetB and PqiB shown in ribbon representation in the context of the periplasm. The position of the peptidoglycan is modeled in. At this position, the distance between ring 4 and 5 in LetB (~37 Å) is considerably larger than the distance between other rings (28.2-32.7 Å), and at this position a poly-proline region introduces a "break" visible in the needle of PqiB. These may accommodate interactions with the peptidoglycan.

**Extended Data Table 1: Data collection and refinement statistics for cryo EM structure of LetB**

| | |
|---|---|
| pixel size (Å): | 1.31 |
| electron energy (kV): | 300 |
| electron dose (e-/Å/px): | 10.0 |
| total electron dose (e-/Å2): | 80.0 |
| number of frames in each movie: | 50 |
| images acquired: | 4906 |
| Number of particles: | |
| picked: | 731,231 |
| selected after 2D classification: | 534,003 |
| Pixel dimension of individual windows: | 280x280 |

| | **Map 1 Model 1** | **Map 2 Model 2** |
|---|---|---|
| Number of particles: | 61,618 | 57,157 |
| Resolution (Å), Relion: | 3.46 | 3.49 |
| Sphericity of 3DFSC: | 0.981 | 0.980 |
| Map CC: | 0.793 | 0.781 |
| rmsd (bonds): | 0.01 | 0.01 |
| rmsd (angles): | 1.28 | 1.18 |
| All-atom clashscore | 3.86 | 3.96 |
| Ramachandran plot values: | | |
| outliers: | 0.00 % | 0.00 % |
| allowed: | 2.71 % | 2.29 % |
| favored: | 97.29 % | 97.71 % |
| Rotamer outliers: | 0.00 % | 0.00 % |
| C-beta deviations: | 0 | 0 |
| Overall score (Molprobity): | 1.31 | 1.25 |
| EMRinger Score: | 3.14 | 2.82 |
| Deposition: | | |
| PDB ID: | TBD | TBD |
| EMDB ID: | TBD | TBD |

| **Composite closed model 9:** | **Map 3 Model 3** | **Map 4 Model 4** | **Map 5 Model 5** |
|---|---|---|---|
| Number of particles: | 101,464 | 144,219 | 147,415 |

| | | | |
|---|---|---|---|
| Resolution (Å), Relion: | 3.06 | 2.96 | 3.03 |
| Sphericity of 3DFSC: | 0.978 | 0.989 | 0.990 |
| Map CC: | 0.796 | 0.800 | 0.806 |
| rmsd (bonds): | 0.01 | 0.01 | 0.01 |
| rmsd (angles): | 1.24 | 1.17 | 1.16 |
| All-atom clashscore | 5.69 | 4.35 | 6.08 |
| Ramachandran plot values: | | | |
| outliers: | 0.00 % | 0.00 % | 0.00 % |
| allowed: | 5.31 % | 1.43 % | 2.40 % |
| favored: | 94.69 % | 98.57 % | 97.60 % |
| Rotamer outliers: | 0.00 % | 0.00 % | 0.00 % |
| C-beta deviations: | 0 | 0 | 0 |
| Overall score (Molprobity): | 1.68 | 1.21 | 1.42 |
| EMRinger Score: | 3.32 | 2.98 | 3.01 |
| Deposition: | | | |
| PDB ID: | TBD | TBD | TBD |
| EMDB ID: | TBD | TBD | TBD |

| **Composite open model 10:** | **Map 6 Model 6** | **Map 7 Model 7** | **Map 8 Model 8** |
|---|---|---|---|
| Number of particles: | 71,235 | 59,687 | 21,149 |
| Resolution (Å), Relion: | 3.60 | 3.43 | 3.78 |
| Sphericity of 3DFSC: | 0.986 | 0.958 | 0.967 |
| Map CC: | 0.722 | 0.789 | 0.814 |
| rmsd (bonds): | 0.01 | 0.01 | 0.01 |
| rmsd (angles): | 1.16 | 1.33 | 1.19 |
| All-atom clashscore | 4.20 | 4.38 | 5.14 |
| Ramachandran plot values: | | | |
| outliers: | 0.00 % | 0.00 % | 0.25 % |
| allowed: | 4.25 % | 1.15 % | 5.76 % |
| favored: | 95.75 % | 98.85 % | 93.99 % |
| Rotamer outliers: | 0.00 % | 0.00 % | 0.00 % |
| C-beta deviations: | 0 | 0 | 0 |
| Overall score (Molprobity): | 1.50 | 1.22 | 1.68 |
| EMRinger Score: | 2.34 | 3.27 | 2.89 |
| Deposition: | | | |
| PDB ID: | TBD | TBD | TBD |
| EMDB ID: | TBD | TBD | TBD |

**Extended Data Table 2: Data collection and refinement statistics of LetB MCE2-MCE3 crystal structure**

| **Crystal structure of MCE2-MCE3** | |
|---|---|
| **Data collection** | |
| Space group: | P65 |
| Cell dimensions: | |
| a, b, c (Å): | 87.56, 87.56, 116.99 |
| α, β, ɣ (°): | 90, 90, 120 |
| Resolution (Å): | 46.32-2.15 (2.21-2.15)[1] |
| Wavelength (Å): | 1.0000 |
| Observations: | 259,635 |
| Unique Reflections: | 27,654 |
| Redundancy: | 9.4 (4.8) |
| Completeness (%): | 99.8 (97.4) |
| CC1/2: | 1.00 (0.62) |
| $I/\sigma I$ | 21.3 (1.2) |
| $R_{meas}$ | 0.05 (1.26) |
| **Refinement** | |
| Resolution (Å): | 41.0 - 2.15 |
| Reflections (work): | 26,145 |
| Reflections (free): | 1,321 |
| $R_{work}$ / $R_{free}$ (%): | 21.4 / 24.4 |
| No. atoms: | |
| Protein: | 3,263 |
| Water: | 48 |
| Other: | 0 |
| Mean B-factor: | |
| Protein: | 84.1 |
| Water: | 61.8 |
| R.M.S. Deviations: | |
| Bond lengths (Å): | 0.003 |
| Bond angles (°): | 0.56 |
| Ramachandran plot: | |
| Favored: | 97.2% |
| Outliers: | 0.2 % |
| Rotamer outliers: | 0.56 % |
| Molprobity: | |
| Molprobity score: | 1.17 |
| Percentile: | 100th |
| All-atom clashscore: | 2.44 |
| Percentile: | 99th |
| PDB ID: | TBD |

[1] Values in parentheses are for highest-resolution shell.

#### Supplementary Table 1: Constructs used in this study

| Plasmid ID | Features | Use in this study |
|---|---|---|
| pBEL1620 | LetAB (from previous study(3)) | WT complementation plasmid |
| pBEL1324 | LetB(43-877)-6xHis | Expression plasmid for WT LetB (without TM region) for cryo and negative stain EM |
| pBEL1584 | LetB(159-383)-6xHis | Expression plasmid of rings 2 and 3 of LetB for crystallography |
| pBEL1751 | Desulfocapsa_sulfexigens_UWK_02604(38-533)-6xHis | Expression plasmid for naturally occurring 4xMCE protein (without TM region) |
| pBEL1752 | Nitrincola_nitratireducens_D791_00999(47-644)-6xHis | Expression plasmid for naturally occurring 5xMCE protein (without TM region) |
| pBEL1753 | Pseudomonas_aeruginosa_PA4689(41-768)-6xHis | Expression plasmid for naturally occurring 6xMCE protein (without TM region) |
| pBEL1755 | Denitrovibrio_acetiphilus_Dacet_0211(41-988)-6xHis | Expression plasmid for naturally occurring 8xMCE protein (without TM region) |
| pBEL2005 | LetA-LetB(Δ161-747) | Complementation plasmid for 2 ring LetB deletion |
| pBEL2006 | LetA-LetB(Δ161-634) | Complementation plasmid for 3 ring LetB deletion |
| pBEL2007 | LetA-LetB(Δ161-514) | Complementation plasmid for 4 ring LetB deletion |
| pBEL2008 | LetA-LetB(Δ515-746) | Complementation plasmid for 5 ring LetB deletion |
| pBEL2009 | LetA-LetB(Δ634-746) | Complementation plasmid for 6 ring LetB deletion |
| pBEL2000 | LetB(43-877; Δ161-747)-6xHis | Expression plasmid for 2 ring LetB deletion (without TM region) |
| pBEL2001 | LetB(43-877; Δ161-634)-6xHis | Expression plasmid for 3 ring LetB deletion (without TM region) |
| pBEL2002 | LetB(43-877; Δ161-514)-6xHis | Expression plasmid for 4 ring LetB deletion (without TM region) |
| pBEL2003 | LetB(43-877; Δ515-746)-6xHis | Expression plasmid for 5 ring LetB deletion (without TM region) |
| pBEL2004 | LetB(43-877; Δ634-746)-6xHis | Expression plasmid for 6 ring LetB deletion (without TM region) |
| pBEL1886 | LetA-LetB(Δ114-128, P113G) | Complementation plasmid for PLL1 deletion |
| pBEL1887 | LetA-LetB(Δ229-239, D228G, L240G) | Complementation plasmid for PLL2 deletion |
| pBEL1888 | LetA-LetB(Δ347-360, P346G) | Complementation plasmid for PLL3 deletion |

| | | |
|---|---|---|
| pBEL1889 | LetA-LetB(Δ459-360, D458G, L469G) | Complementation plasmid for PLL4 deletion |
| pBEL1890 | LetA-LetB(Δ581-590, K580G, Q591G) | Complementation plasmid for PLL5 deletion |
| pBEL1891 | LetA-LetB(Δ703-708, Q702G, E709G) | Complementation plasmid for PLL6 deletion |
| pBEL1892 | LetA-LetB(Δ816-826, S815G, T827G) | Complementation plasmid for PLL7 deletion |
| pBEL1993 | LetB(43-877; Δ114-128, P113G)-6xHis | Expression plasmid for LetB PLL1 deletion (without TM region) |
| pBEL1994 | LetB(43-877; Δ229-239, D228G, L240G)-6xHis | Expression plasmid for LetB PLL2 deletion (without TM region) |
| pBEL1995 | LetB(43-877; Δ347-360, P346G)-6xHis | Expression plasmid for LetB PLL3 deletion (without TM region) |
| pBEL1996 | LetB(43-877; Δ459-360, D458G, L469G)-6xHis | Expression plasmid for LetB PLL4 deletion (without TM region) |
| pBEL1997 | LetB(43-877; Δ581-590, K580G, Q591G)-6xHis | Expression plasmid for LetB PLL5 deletion (without TM region) |
| pBEL1998 | LetB(43-877; Δ703-708, Q702G, E709G)-6xHis | Expression plasmid for LetB PLL6 deletion (without TM region) |
| pBEL1999 | LetB(43-877; Δ816-826, S815G, T827G)-6xHis | Expression plasmid for LetB PLL7 deletion (without TM region) |
| pBEL1857 | LetA-LetB(L243N) | Complementation plasmid for LetB mutant L243N |
| pBEL1858 | LetA-LetB(L246N) | Complementation plasmid for LetB mutant L246N |
| pBEL1859 | LetA-LetB(V247N) | Complementation plasmid for LetB mutant V247N |
| pBEL1860 | LetA-LetB(L355N) | Complementation plasmid for LetB mutant L355N |
| pBEL1861 | LetA-LetB(L358N) | Complementation plasmid for LetB mutant L358N |
| pBEL1862 | LetA-LetB(L359N) | Complementation plasmid for LetB mutant L359N |
| pBEL1806 | LetB(43-877; L243N)-6xHis | Expression plasmid for LetB mutant L243N |
| pBEL1807 | LetB(43-877; L246N)-6xHis | Expression plasmid for LetB mutant L246N |
| pBEL1808 | LetB(43-877; V247N)-6xHis | Expression plasmid for LetB mutant V247N |
| pBEL1809 | LetB(43-877; L355N)-6xHis | Expression plasmid for LetB mutant L355N |
| pBEL1810 | LetB(43-877; L358N)-6xHis | Expression plasmid for LetB mutant L358N |
| pBEL1811 | LetB(43-877; L359N)-6xHis | Expression plasmid for LetB mutant L358N |
| pBEL1725 | LetB(43-877; F468Amber)-6xHis | Expression plasmid for LetB mutant F468Bpa |
| pBEL1726 | LetB(43-877; W476Amber)-6xHis | Expression plasmid for LetB mutant W476pa |
| pBEL1727 | LetB(43-877; Y814Amber)-6xHis | Expression plasmid for LetB mutant Y814Bpa |

| | | |
|---|---|---|
| pBEL1728 | LetB(43-877; F818Amber)-6xHis | Expression plasmid for LetB mutant F818Bpa |
| pBEL1729 | LetB(43-877; F833Amber)-6xHis | Expression plasmid for LetB mutant F833Bpa |
| pBEL1730 | LetB(43-877; K488Amber)-6xHis | Expression plasmid for LetB mutant K488Bpa |
| pBEL1731 | LetB(43-877; E854Amber)-6xHis | Expression plasmid for LetB mutant E854Bpa |

**Supplementary Video 1.** First three eigenvectors obtained from the multi-body refinement analysis of Map1 in Relion 3.0, and using the 3D variability analysis in Cryosparc 2.9. This analysis shows possible rotation of the rings relative to each other, as well as possible bending and elongation of LetB.

**Supplementary Video 2.** Morph between the open and closed states of LetB. These models are generated from composite open and closed maps.

**Supplementary Video 3.** Morph between the open and closed states of PqiB.

### Supplementary Data Figure 1

**Supplementary Data Figure 1. Gels for detection of photocrosslinked substrate in tunnel of LetB.** Uncropped gels corresponding to **Fig. 5h**.

**a –** replicate 1 (uncropped versions of gels in main figure 5c)

##### crosslinked

WT F468B W476B Y814B F818B F833B K488B E854B

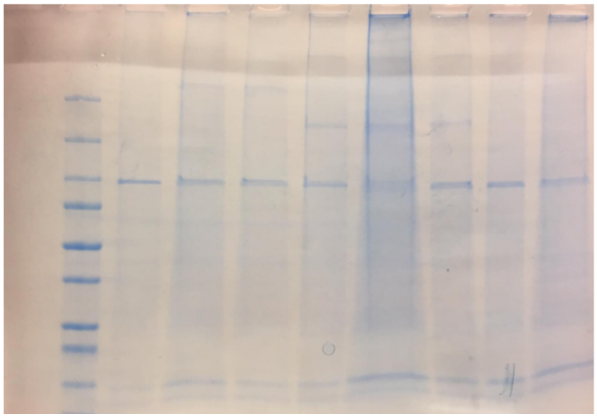

coomassie

WT F468B W476B Y814B F818B F833B K488B E854B

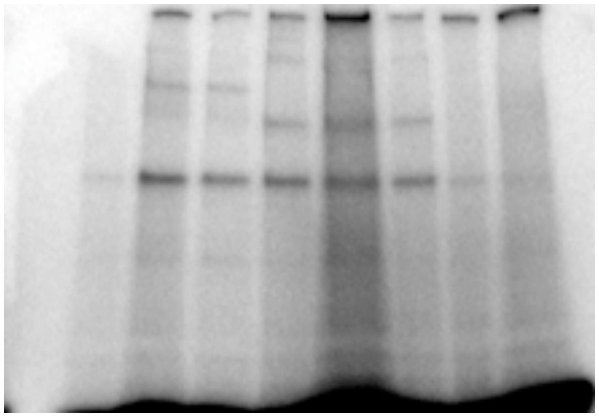

phosphor-screen

##### uncrosslinked

WT F468B W476B Y814B F818B F833B K488B E854B

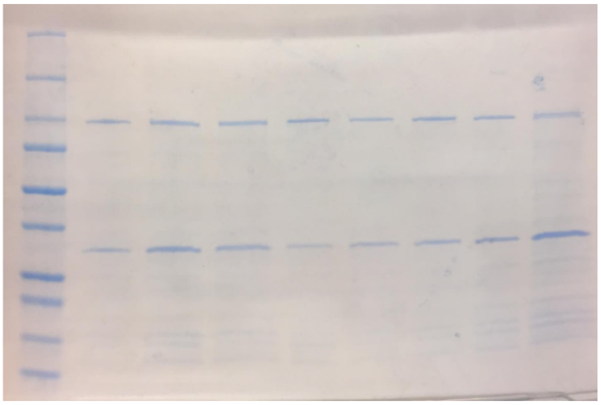

coomassie

WT F468B W476B Y814B F818B F833B K488B E854B

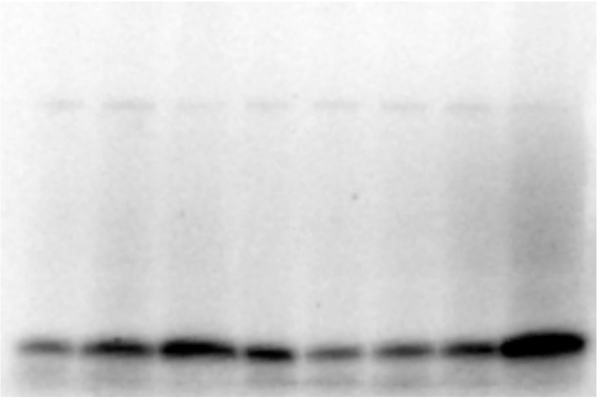

phosphor-screen

#### Supplementary Data Figure 2

**Supplementary Data Fig. 2. Representative micrographs and 2D class averages for all negative stain EM data.**

**a –** Naturally occurring MCE proteins with 4, 5, 6 and 8 MCE domains from other species raw micrographs

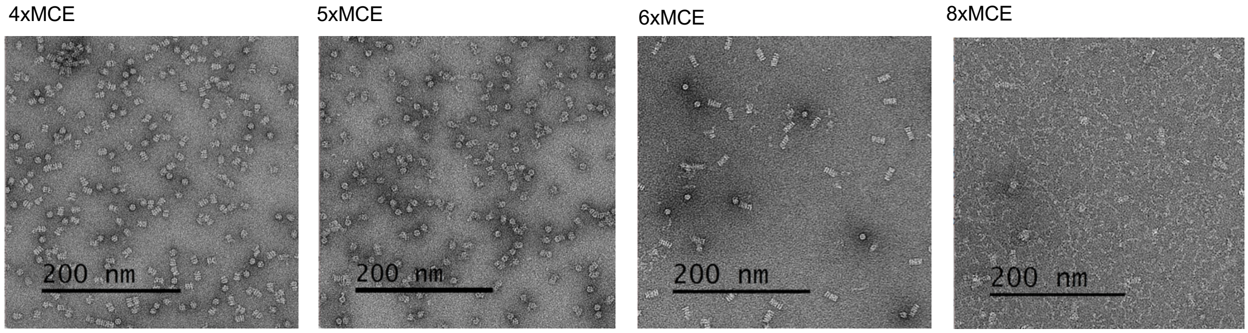


**b –** Naturally occurring MCE proteins with 4, 5, 6 and 8 MCE domains from other species 2D classes

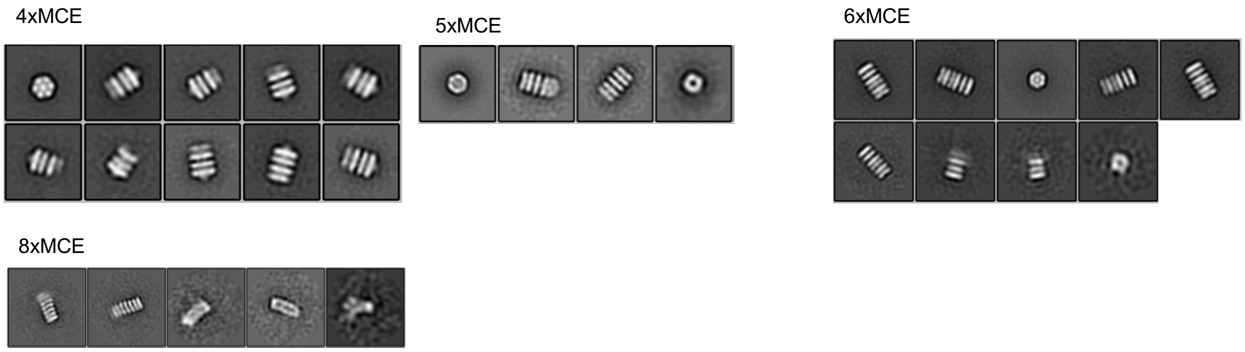

### Supplementary Data Figure 2 (cont)

**c –** LetB truncations raw micrographs

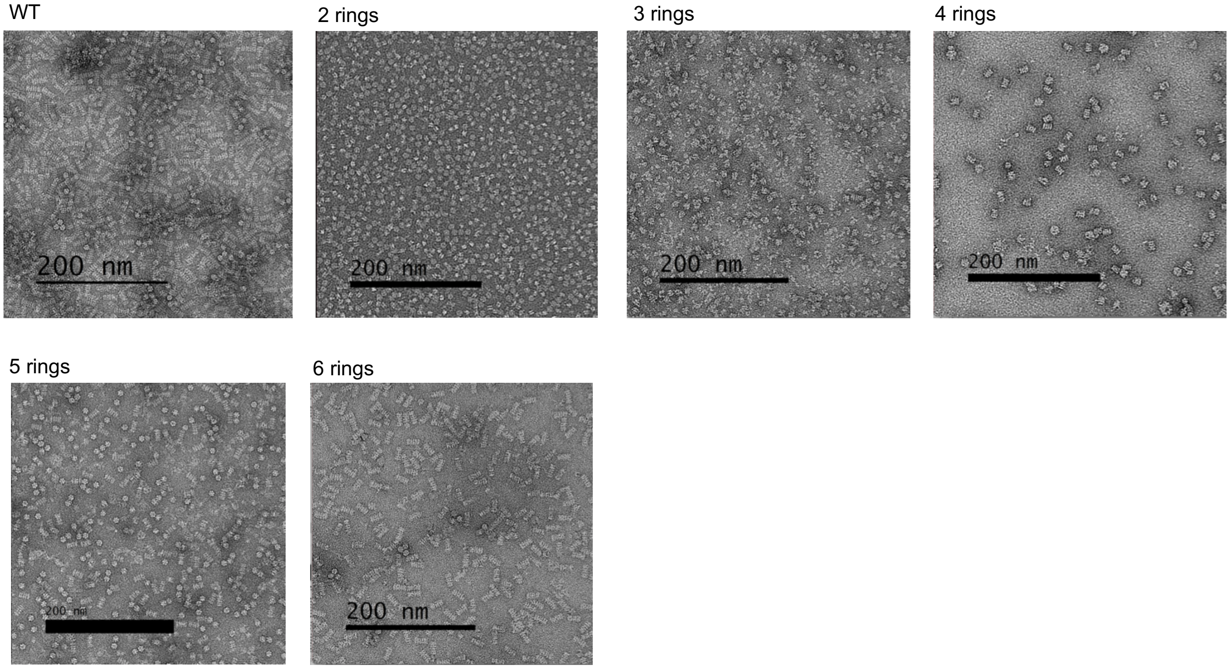


**d –** LetB truncations 2D classes

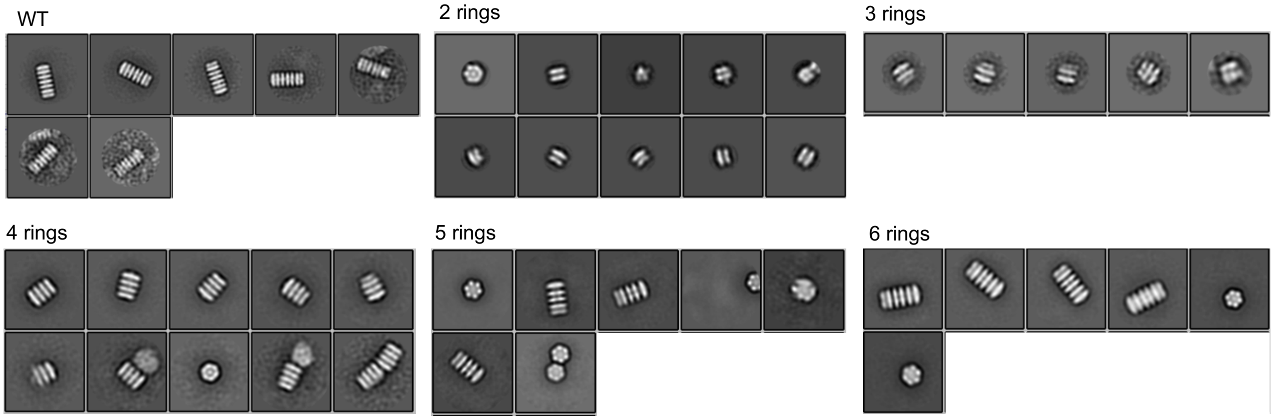

#### Supplementary Data Figure 2 (cont)

**e –** Pore lining loop deletions raw micrographs

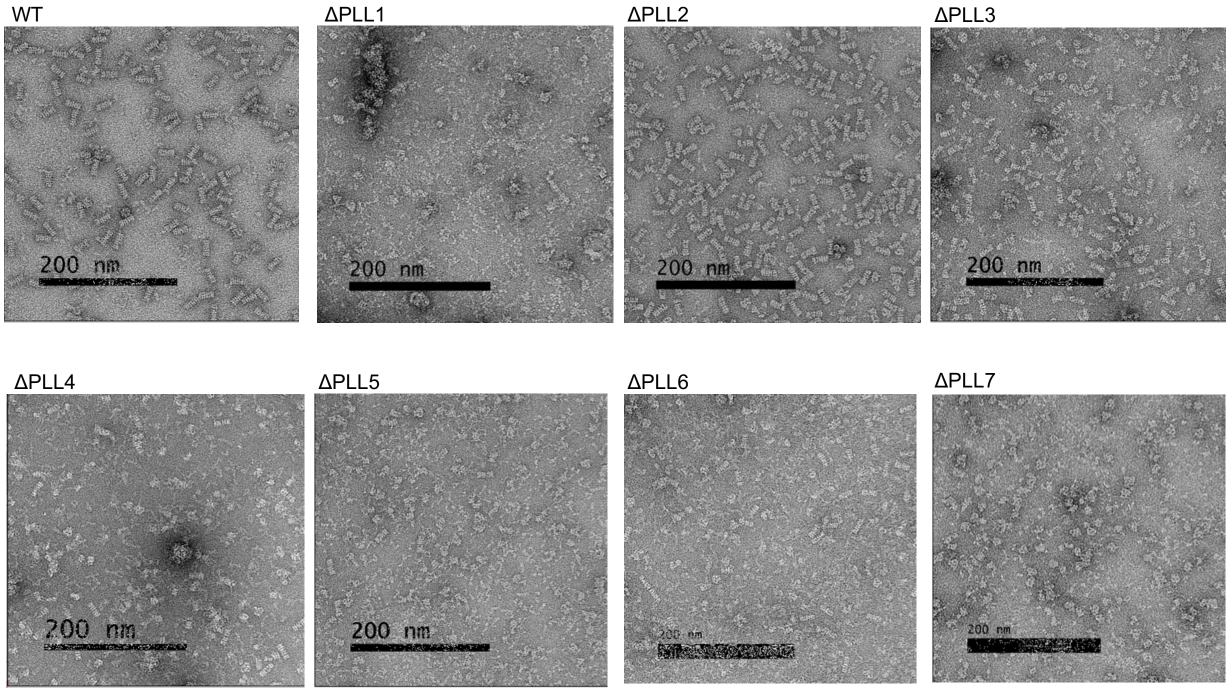


**f –** Pore lining loop deletions 2D classes

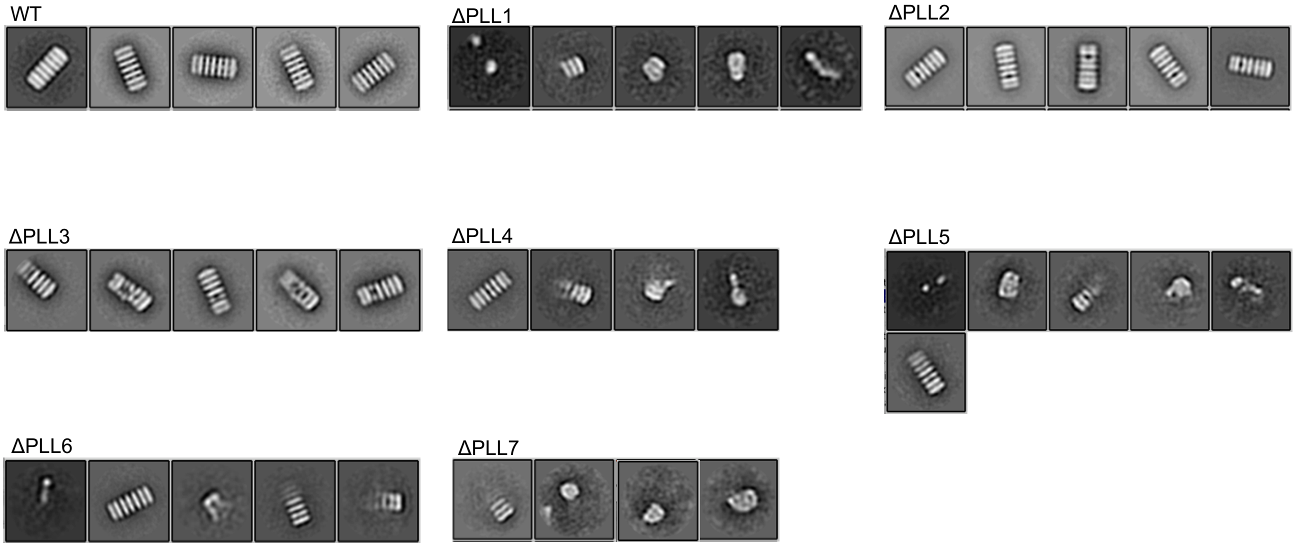

### Supplementary Data Figure 2 (cont)

**g –** Hydrophobic motif mutants raw micrographs

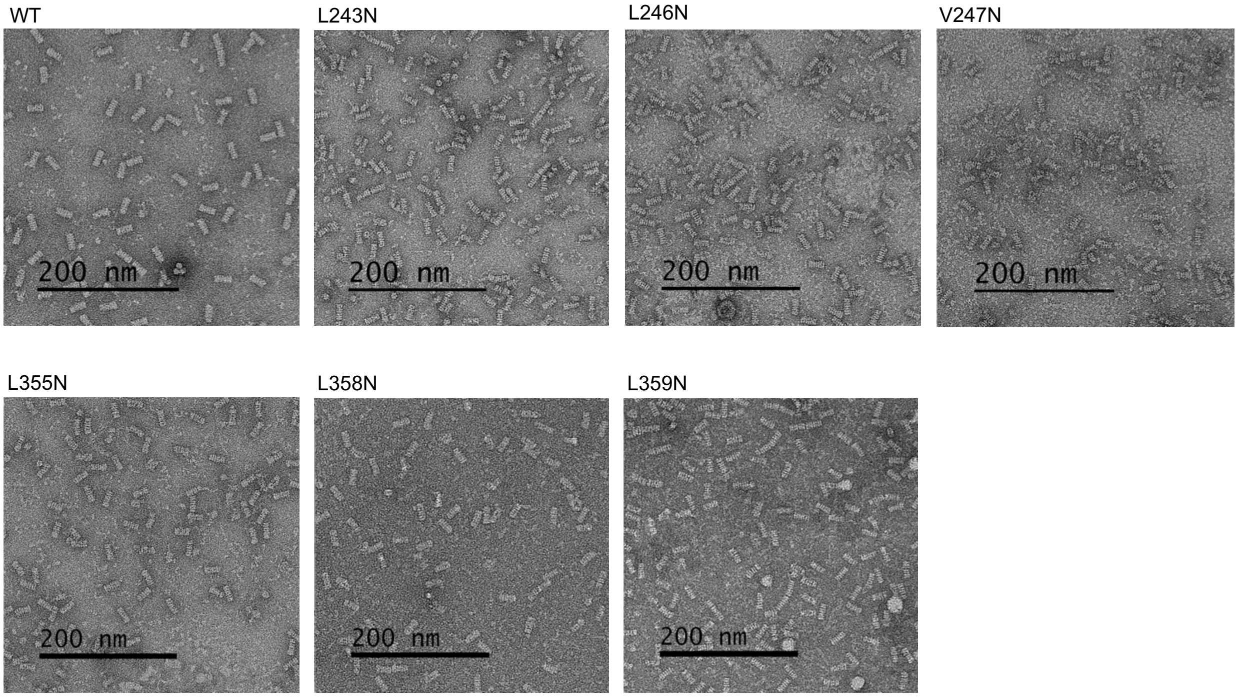


**h –** Hydrophobic motif mutants 2D classes

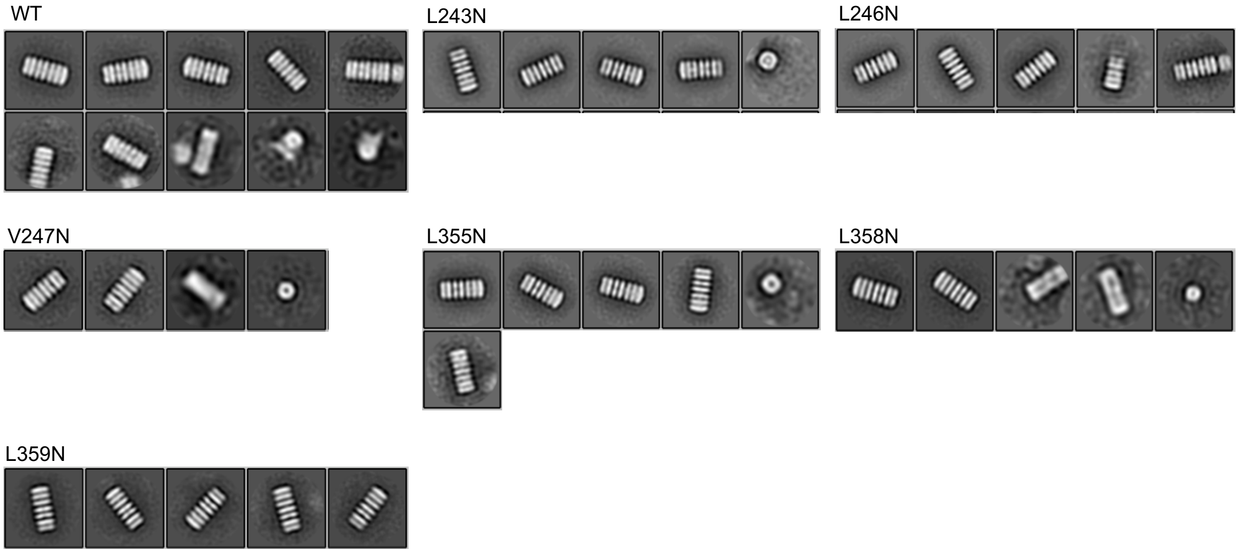
